# Antibody-mediated broad sarbecovirus neutralization through ACE2 molecular mimicry

**DOI:** 10.1101/2021.10.13.464254

**Authors:** Young-Jun Park, Anna De Marco, Tyler N Starr, Zhuoming Liu, Dora Pinto, Alexandra C. Walls, Fabrizia Zatta, Samantha K. Zepeda, John Bowen, Kaitlin S Sprouse, Anshu Joshi, Martina Giurdanella, Barbara Guarino, Julia Noack, Rana Abdelnabi, Shi-Yan Caroline Foo, Florian A. Lempp, Fabio Benigni, Gyorgy Snell, Johan Neyts, Sean PJ Whelan, Herbert W. Virgin, Jesse D Bloom, Davide Corti, Matteo Samuele Pizzuto, David Veesler

**Affiliations:** Department of Biochemistry, University of Washington, Seattle, WA 98195, USA; Humabs Biomed SA, a subsidiary of Vir Biotechnology, 6500 Bellinzona, Switzerland; Basic Sciences Division, Fred Hutchinson Cancer Research Center, Seattle, WA 98109, USA; Department of Molecular Microbiology, Washington University School of Medicine, St. Louis, MO, 63110, USA; Vir Biotechnology, San Francisco, CA, USA; Rega Institute for Medical Research, Laboratory of Virology and Chemotherapy, KU Leuven, Belgium; Howard Hughes Medical Institute, Seattle, WA 98109

## Abstract

Understanding broadly neutralizing sarbecovirus antibody responses is key to developing countermeasures effective against SARS-CoV-2 variants and future spillovers of other sarbecoviruses. Here we describe the isolation and characterization of a human monoclonal antibody, designated S2K146, broadly neutralizing viruses belonging to all three sarbecovirus clades known to utilize ACE2 as entry receptor and protecting therapeutically against SARS-CoV-2 beta challenge in hamsters. Structural and functional studies show that most of the S2K146 epitope residues are shared with the ACE2 binding site and that the antibody inhibits receptor attachment competitively. Viral passaging experiments underscore an unusually high barrier for emergence of escape mutants making it an ideal candidate for clinical development. These findings unveil a key site of vulnerability for the development of a next generation of vaccines eliciting broad sarbecovirus immunity.

---

The zoonotic spillover of SARS-CoV-2 has resulted in a global pandemic causing over 231 million infections and more than 4.7 million fatalities as of September 2021. Continued SARS-CoV-2 evolution led to the emergence of variants of concern (VOC) that are characterized by higher transmissibility, immune evasion and/or disease severity. There is therefore a need for developing pan-sarbecovirus countermeasures, such as vaccines and therapeutics that are effective against all SARS-CoV-2 variants and divergent zoonotic sarbecoviruses for pandemic preparedness (*1*).

The coronavirus spike glycoprotein (S) promotes viral entry into host cells and is the main target of neutralizing antibodies elicited by infection or vaccination (*2–6*). S comprises an S_1_ subunit, which recognizes host cell receptors, and an S_2_ subunit that drives viral-cell membrane fusion. The S_1_ subunit includes the N-terminal domain and the receptor-binding domain (RBD), the latter domain interacts with angiotensin-converting enzyme 2 (ACE2) to allow virus entry into host cells in the case of SARS-CoV and SARS-CoV-2 (*4, 7–10*). The RBD is the target of the majority of serum neutralizing activity elicited by infection (*11*) and vaccination (*12*) and exposes multiple antigenic sites in the domain core that are recognized by broadly neutralizing sarbecovirus antibodies (Abs) (*13–19*). However, a large fraction of Abs in polyclonal sera (*11*) and most monoclonal Abs (mAbs) selected for therapeutic development (*20*) target epitopes overlapping the ACE2-contact surface (designated the receptor-binding motif (RBM)). The marked genetic divergence and plasticity of the RBM among SARS-CoV-2 variants and sarbecoviruses limits the breadth of Abs recognizing this region and they are readily escaped by mutations (*14, 21–26*).

## Identification and characterization of the S2K146 mAb

To identify broadly neutralizing sarbecovirus Abs, we isolated SARS-CoV-2 S-specific (IgG) memory B cells from one symptomatic COVID-19 convalescent individual (who was not hospitalized) 35 days after symptoms onset. We identified one mAb, designated S2K146 (IGHV3-43; IGL1-44), which did not compete with either S309 (site IV) (*15*) or S2×259 (site II) (*13*) for binding to the SARS-CoV-2 RBD but competed with the RBM-targeted S2E12 mAb (site I) (*27*) **(Fig 1A)**. Like S2E12, S2K146 bound to all SARS-CoV-2 VOC RBDs as well as all clade 1b sarbecovirus RBDs tested by ELISA **(Fig 1B and Fig S1A)**. In contrast to S2E12 and other site I-targeting Abs described so far, S2K146 also cross-reacted with SARS-CoV and WIV-1 RBDs (clade 1a), which share 73% and 76% sequence identity with the SARS-CoV-2 RBD, respectively **(Fig 1B and Fig S1A)**. S2K146 did not bind to clades 2 and 3 sarbecovirus RBDs, similarly to the broadly neutralizing sarbecovirus S309 mAb but in contrast to the S2×259 or S2H97 mAbs (*13, 14*). Consistent with this finding we observed a similar pattern of S2K146 cross-reactivity with clades 1a and 1b sarbecoviruses using native S trimers transiently expressed on the surface of mammalian cells **(Fig 1C)** and yeast-surface displayed RBDs **(Fig 1D)**. Based on these results, we hypothesized that S2K146 recognizes a previously uncharacterized RBM epitope which is conserved among sarbecovirus clades 1a and 1b.

**Figure 1.**
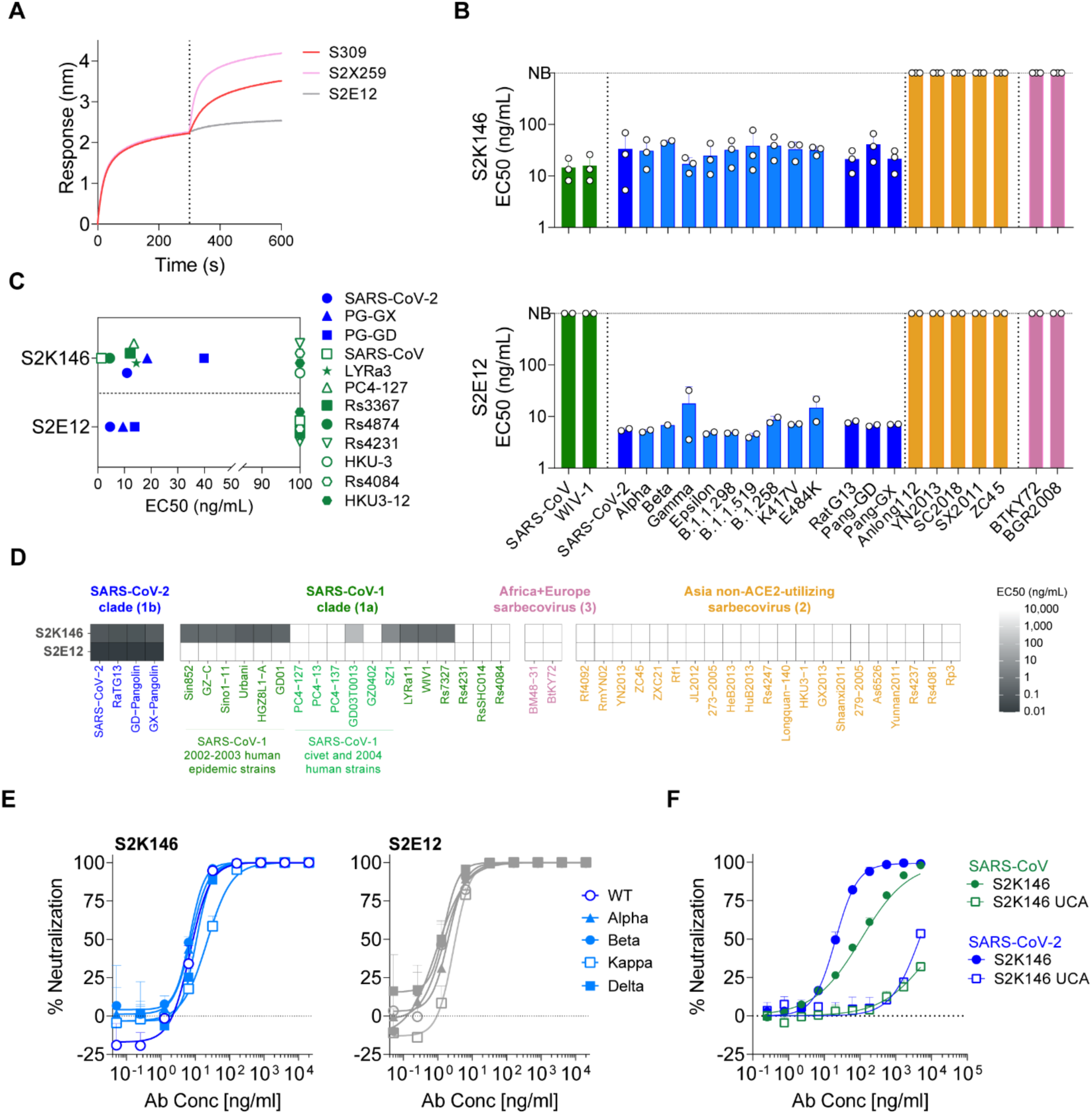
Identification of the S2K146 cross-reactive and broadly neutralizing sarbecovirus mAb. **A**) Binding of site I-targeting S2E12, site IV-targeting S309, or site II-targeting S2×259 recombinant IgG1 (second phase) following association of S2K146 mAb (first phase) to His-tagged SARS-CoV-2 RBD immobilized on anti-His sensors, as measured by biolayer interferometry. **B)** Cross-reactivity of S2K146 (upper panel) and S2E12 (lower panel) with 22 RBDs from sarbecovirus clades (in brackets) and SARS-CoV-2 variants analyzed by ELISA. EC50 of at least two independent experiments are shown. **C)** Flow cytometry analysis of S2K146 cross-reactivity with a panel of 12 S glycoproteins representative of sarbecovirus clades 1a and 1b transiently expressed on the surface of mammalian cells. **D)** S2K146 cross-reactivity with sarbecovirus RBDs displayed at the surface of yeast. **E**) S2K146- and S2E12-mediated neutralization of replication-competent SARS-CoV-2 (USA-WA1/2020) and SARS-CoV-2 VOC viruses. **F**) S2K146 and S2K146-UCA-mediated neutralization of VSV pseudotypes harboring SARS-CoV-2 S or SARS-CoV S.

To evaluate the neutralization potency of the S2K146 mAb, we carried out dose-response inhibition assays using a vesicular stomatitis virus (VSV) pseudotyping platform. S2K146 efficiently blocked SARS-CoV S- and SARS-CoV-2 S-mediated entry into cells with half maximum inhibitory concentration (IC_50_) of 108 and 16 ng/mL, respectively **(Fig S1B)**. Moreover, S2K146 potently neutralized VSV pseudotypes harboring SARS-CoV-2 S glycoproteins from VOC including Alpha, Beta, Gamma, Delta plus-AY.1/AY.2, Epsilon and Lambda **(Fig S1B)**. Moreover, S2K146 neutralized weakly VSV pseudotyped with BtKY72 S (clade 3) harboring the K493Y/T498W S mutations (SARS-CoV-2 numbering) (*28*), which allow ACE2 mediated entry in mammalian cells (**Fig S1C**). Finally, S2K146 neutralized authentic SARS-CoV-2 (isolate USA-WA1/2020, lineage A, IC_50_= 10 ng/mL) and SARS-CoV-2 VOC (Alpha, IC_50_= 9 ng/mL; Beta, IC_50_= 9 ng/mL; Delta, IC_50_= 8 ng/mL; Kappa, IC_50_= 30 ng/mL) with a potency approaching that observed with the ultrapotent S2E12 mAb (*27*) (Wuhan-1, IC_50_= 3.5 ng/mL; Alpha, IC_50_= 2.5 ng/mL; Beta, IC_50_= 2 ng/mL; Delta, IC_50_= 1.5 ng/mL; Kappa, IC_50_= 4.5 ng/mL) **(Fig 1E)**.

To assess the role of somatic mutations for S2K146 binding and neutralization, we generated its inferred unmutated common ancestor (S2K146 UCA). Alignment with the UCA amino acid sequence reveals that S2K146 harbors seven and two somatic hypermutations in the heavy- and light-chain complementarity determining regions (CDR), respectively (VH identity: 94.4% and VL identity: 98.9%, **Fig. S1D)**. Except for WIV1, no differences between S2K146 and S2K146 UCA were observed in terms of cross-reactivity to a panel of RBDs representative of clade 1 sarbecoviruses, as determined by ELISA **(Fig S1E)**. Nevertheless, biolayer interferometry revealed that S2K146 bound to SARS-CoV and SARS-CoV-2 prefusion-stabilized S ectodomain trimers with enhanced avidities compared to S2K146 UCA **(Fig S1F)**. Accordingly, S2K146 UCA showed a marked loss in neutralizing activity against both SARS-CoV S and SARS-CoV-2 S VSV pseudotypes **(Fig 1F)**. Our results suggest that somatic hypermutations associated with S2K146 affinity maturation are important to enhance antibody avidity and potency but not necessarily breadth.

These data establish S2K146 as a potent cross-reactive mAb which neutralizes SARS-CoV-2- and SARS-CoV-related sarbecoviruses as well as circulating VOC via recognition of an antigenic site distinct from previously identified broadly neutralizing mAbs (*13–19*). Our data further show that S2K146 neutralized a clade 3 sarbecovirus (BtKY72) with two RBD mutations enabling utilization of human ACE2 as entry receptor, suggesting that this mAb could also inhibit other clade 3 bat viruses that would acquire similar adaptive mutations.

## Structural basis for S2K146-mediated broad sarbecovirus neutralization

To understand the unique sarbecovirus cross-reactivity of the RBM-specific S2K146 mAb, we determined a cryo-electron microscopy structure of the S2K146 Fab fragment in complex with the SARS-CoV-2 S ectodomain trimer at 3.2 Å resolution **(Fig 2A, Fig S2 and Table S1)**. 3D classification of the data led to the determination of a structure with three open RBDs each bound to a S2K146 Fab as well as a structure with two open RBDs and one closed RBD with a Fab bound to each of them **(Fig S2)**. Our cryoEM data show that opening of two RBDs is enough to allow three Fabs to bind to an S trimer, as the remaining closed RBD can engage an S2K146 Fab due to its angle of approach.

**Figure 2.**
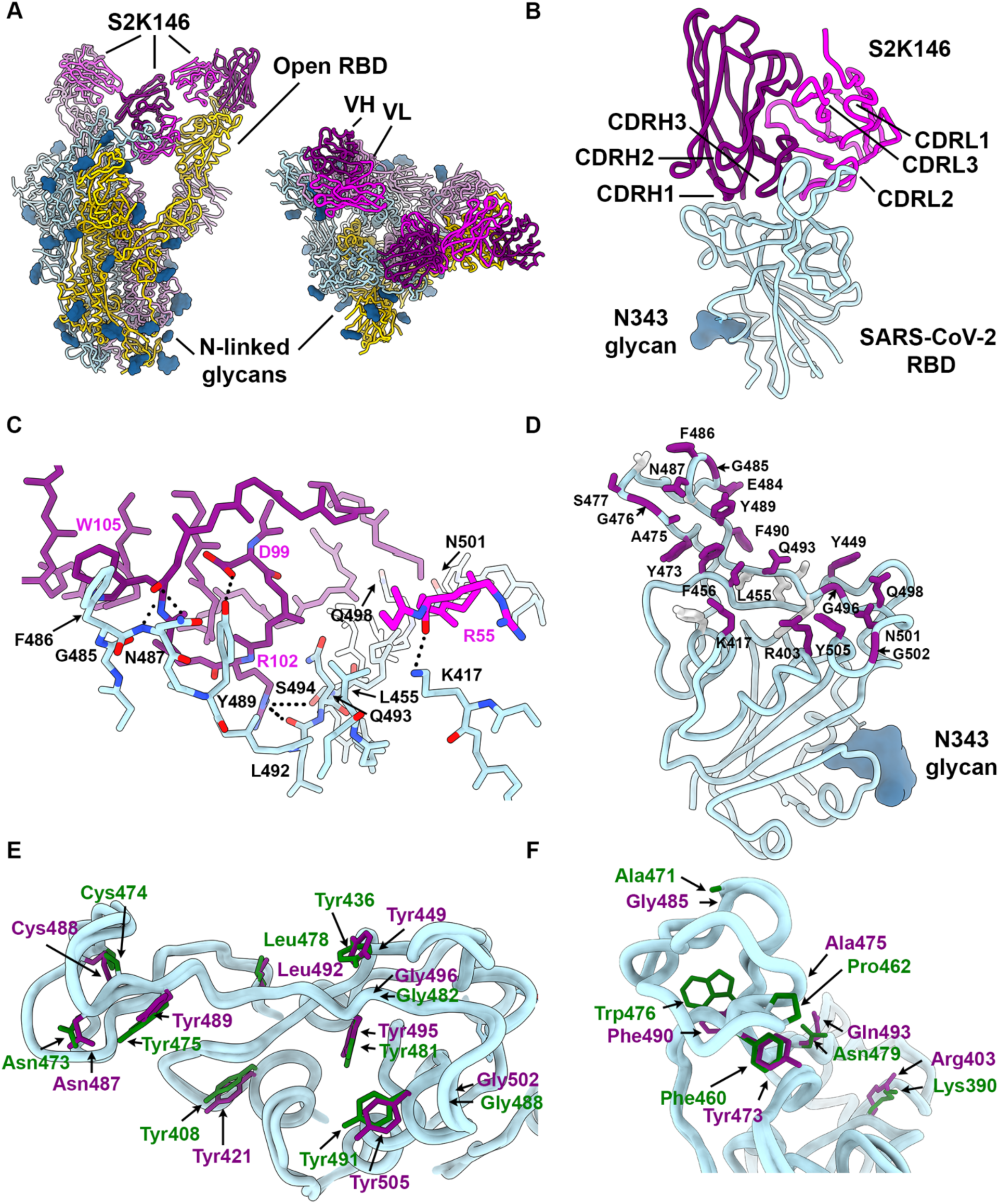
The S2K146 broadly neutralizing mAb recognizes RBD antigenic site I. **A)** CryoEM structure of the prefusion SARS-CoV-2 S ectodomain trimer with three S2K146 Fab fragments bound to two open RBDs and one partially closed RBD viewed along two orthogonal orientations. SARS-CoV-2 S protomers are colored cyan, pink and gold. S2K146 heavy chain and light variable domains are colored purple and magenta. Glycans are rendered as blue spheres. **B)** Ribbon diagram of the S2K146-bound SARS-CoV-2 RBD. **C)** Zoomed-in view of the contacts formed between S2K146 and the SARS-CoV-2 RBD. Selected epitope residues are shown as sticks and electrostatic interactions are indicated with dashed lines. **D)** S2K146 epitope residues shown as sticks and colored purple (labeled) if they are involved in ACE2 binding or grey otherwise (unlabeled). **E)** The side chains of the ten S2K146 epitope residues conserved between the SARS-CoV-2 (purple) and SARS-CoV (PDB 2AJF (*29*), green) RBDs are shown as sticks. **F)** The side chains of the six S2K146 epitope residues conservatively substituted between the SARS-CoV-2 (purple) and SARS-CoV (green) RBDs are shown as sticks.

To overcome the conformational heterogeneity of the S2K146-bound RBDs relative to the rest of the S trimer, we used focused 3D classification and local refinement of the S2K146 variable domains and RBD to obtain a reconstruction at 3.2 Å resolution enabling unambiguous model building and providing a detailed view of the binding interface **(Fig 2B, Fig S2 and Table S1)**. S2K146 recognizes an epitope in antigenic site Ia (*11*), which overlaps with the RBM and is masked when the three RBDs adopt a closed state **(Fig 2A-B)**. The S2K146 paratope includes the heavy chain N-terminus and CDRH1, H2 and H3, accounting for 2/3 of the surface buried upon binding, and light chain CDRL1, L2 and L3, making up the rest of the interface. A total of 1,000 Å^2^ of the paratope surface is buried at the interface with the RBM through hydrogen bonding and shape complementarity.

The S2K146 footprint on the SARS-CoV-2 RBD strikingly resembles that of the ACE2 receptor with 21 out of 28 epitope residues shared with the ACE2 binding site, including key ACE2-contact positions L455, F486, Q493, Q498 and N501 **(Fig 2C-D)**. Moreover, electrostatic interactions formed between S2K146 and the SARS-CoV-2 RBD recapitulate some of the hydrogen bonds or salt bridges involved in ACE2 binding, such as with residues K417, N487 and Y489 **(Fig 2D)**. Although S2K146 contact residues are mutated in several variants, such as K417 (Beta and Gamma), L452 (Delta, Epsilon and Kappa), E484 (Beta, Gamma and Kappa) and N501 (Alpha, Beta and Gamma), the binding interface is resilient to these residue substitutions, in line with retention of binding and neutralization of these variants **(Fig 1E and Fig S1B)**. The cross-reactivity with and broad neutralization of SARS-CoV by S2K146 is explained by the strict conservation or conservative substitution of ten and six epitope residues, respectively **(Fig 2E-F and Fig S3)**, consistent with the ability of each of these RBDs to bind human ACE2.

S2K146 therefore defines an unexpected class of broadly neutralizing mAb engaging the RBM and overcoming mutations found in SARS-CoV-2 variants through molecular mimicry of the ACE2 receptor. As S2K146 does not compete with broadly neutralizing sarbecovirus mAbs targeting other antigenic sites, such as S309 (*15*) and S2×259 (*13*) **(Fig S4)**, they could be combined in a cocktail to enhance breadth further and set an even higher barrier for emergence of escape mutants.

## S2K146 is resilient to a broad spectrum of mutations

To prospectively evaluate the impact of antigenic drift on S2K146 neutralization, we mapped RBD mutations that affect mAb binding using deep-mutational scanning (DMS) of a yeast-displayed RBD mutant library covering all possible single residue substitutions in the Wuhan-Hu-1 RBD background (*24*). S2K146 binding was reduced by only a restricted number of amino acid substitutions, compared to S2E12 which binds an overlapping but distinct epitope **(Fig 3A-D and Fig S5A-B)**. Although mutations at several residues were identified to reduce S2K146 binding by DMS, all of them correspond to RBD residues buried upon ACE2 recognition (F456, A475, E484, F486, N487 and Y489) **(Fig 3B)**. Only one of these residue substitutions (Y489H) is accessible via a single-nucleotide change and could escape S2K146 recognition with a penalty on ACE2 binding affinity smaller than an order of magnitude, as determined by DMS data (*24*).

**Figure 3.**
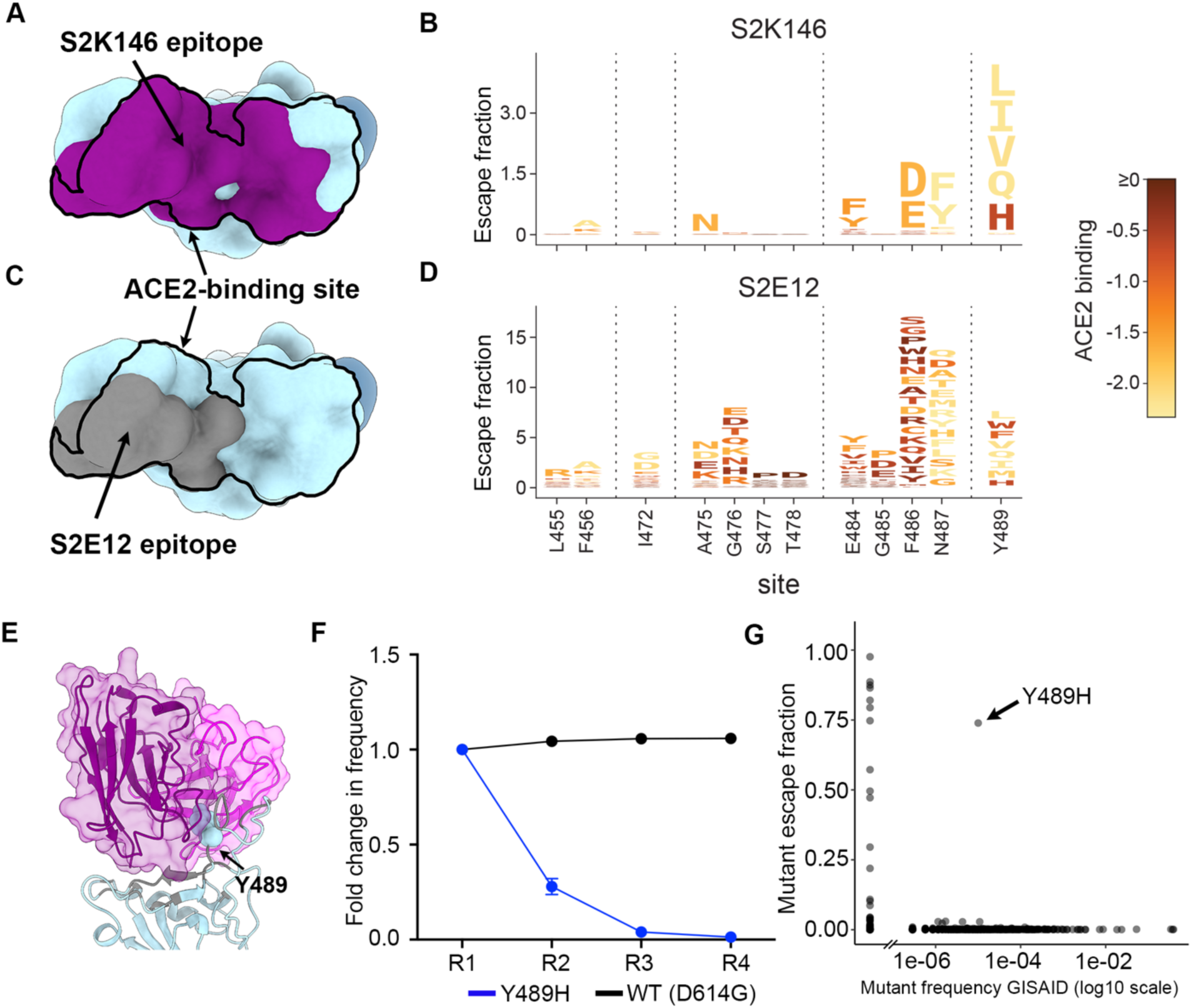
S2K146 is resilient to a broad spectrum of escape mutations. **A)** Molecular surface representation of the SARS-CoV-2 RBD with the S2K146 epitope colored purple and the ACE2 footprint indicated as a black outline. **B)** Mapping of RBD mutations reducing S2K146 binding using DMS of the yeast-displayed SARS-CoV-2 RBD. Sites of strong escape (purple underlines in Supplemental Figure S5.A) are shown in logo plot. Letters are colored according to how mutations affect the RBD affinity for ACE2 as measured via yeast display as measured in Starr et al. (*24*). **C)** Molecular surface representation of the SARS-CoV-2 RBD with the S2E12 epitope colored grey and the ACE2 footprint indicated as a black outline. The N343 glycan is rendered as blue spheres in A and C. **D)** Mapping of RBD mutations reducing S2E12 binding using DMS of the yeast-displayed SARS-CoV-2 RBD. Sites of strong escape (purple underlines in Supplemental Figure S5A) are shown in logo plot, as measured previously in Starr *et. al*(*14*)). **E)** Zoomed-in view of the Y489H S2K146 neutralization escape mutation. The S2K146 heavy and light chain variable domains are shown as transparent purple and magenta surfaces with ribbons, respectively. **F)** Viral replication competition between VSV chimeras harboring the SARS-CoV-2 Wuhan-Hu-1/D614G S with or without the Y489H substitution using VeroE6 cells. **G)** Mutations reducing binding of S2K146 to the RBD based on DMS (escape score) are plotted versus their frequencies among the human-derived SARS-CoV-2 sequences on GISAID as of Sep 27 2021. The large escape mutant (> 5x global median escape fraction) with non-zero frequency is indicated.

To explore whether our escape map was consistent with in vitro viral evolution under mAb pressure, a replication competent VSV-SARS-CoV-2 S Wuhan-Hu-1/D614G chimera (*30*) was grown in cell culture in the presence of the S2K146 mAb. Consistently with the DMS data, Y489H was the sole mutation resulting from a single nucleotide substitution that was detected in all the 36 neutralization-resistant plaques sampled **(Fig 3E, Fig S6 and Table S2)**. SARS-CoV-2 residue Y489 forms multiple interactions with S2K146 CDRH3 and account for 12% of the total epitope buried surface area **(Fig 3E)**, in line with the major impact of the Y489H substitution on mAb neutralization. Out of all the mutations at position 489 identified by DMS to reduce S2K146 binding **(Fig 3B)**, the Y to H substitution had the lowest impact on ACE2 binding, which might explain why it was the sole neutralization escape mutant retrieved upon passaging.

To evaluate the fitness of the Y489H mutant, we carried out a competition assay in which replicating VSV chimeras harboring the Wuhan-Hu-1/D614G S with or without the Y489H substitution were mixed at equal titers and passaged together without mAb. Due to the fitness cost associated with the mutation, the SARS-CoV-2 Y489H S chimera was outcompeted by the Wuhan-Hu-1/D614G chimera after only four rounds of passaging **(Fig 3F)**. Accordingly, only 29 out 2.9 million genomes were found to harbor the S Y489H mutation, underscoring the rarity of and the fitness cost imposed by this residue substitution **(Fig 3G)**. Collectively, these data illustrate the high barrier for emergence of escape mutants imposed by the S2K146 mAb, making it an ideal candidate for clinical development.

## SS2K146 inhibits ACE2 engagement and protects hamsters from SARS-CoV-2 challenge

Our cryoEM structure revealed that S2K146 targets antigenic site Ia which overlaps with the RBM, indicating that mAb binding would compete with ACE2 attachment to the RBD via steric hindrance **(Fig 4A)**. Accordingly, we found that S2K146 inhibited binding of the SARS-CoV-2 and SARS-CoV RBDs to human ACE2 in a concentration-dependent manner, as measured by competition ELISA **(Fig 4B)**. As S2K146 conformationally selects for open RBDs, we assessed if the mAb could promote shedding of the S_1_ subunit from cell-surface-expressed full-length SARS-CoV-2 S, similar to some other RBD-specific mAbs (*11, 13, 18, 27*). S2K146 induced shedding of the S_1_ subunit as efficiently as the RBM-targeting S2E12 mAb, whereas the control mAb S2M11 did not as it locks S in the prefusion closed state (*13*) **(Fig 4C)**. Furthermore, we found that the S2K146 Fab triggered the fusogenic rearrangement of a wildtype-like S ectodomain trimer, as we previously described for several SARS-CoV and SARS-CoV-2 neutralizing mAbs (*14, 31–33*) **(Fig 4D)**. These data show that S2K146-mediated sarbecovirus neutralization relies on competitively blocking viral attachment to the ACE2 receptor and inactivation of S trimers at the surface of virions before encountering host cells.

**Figure 4.**
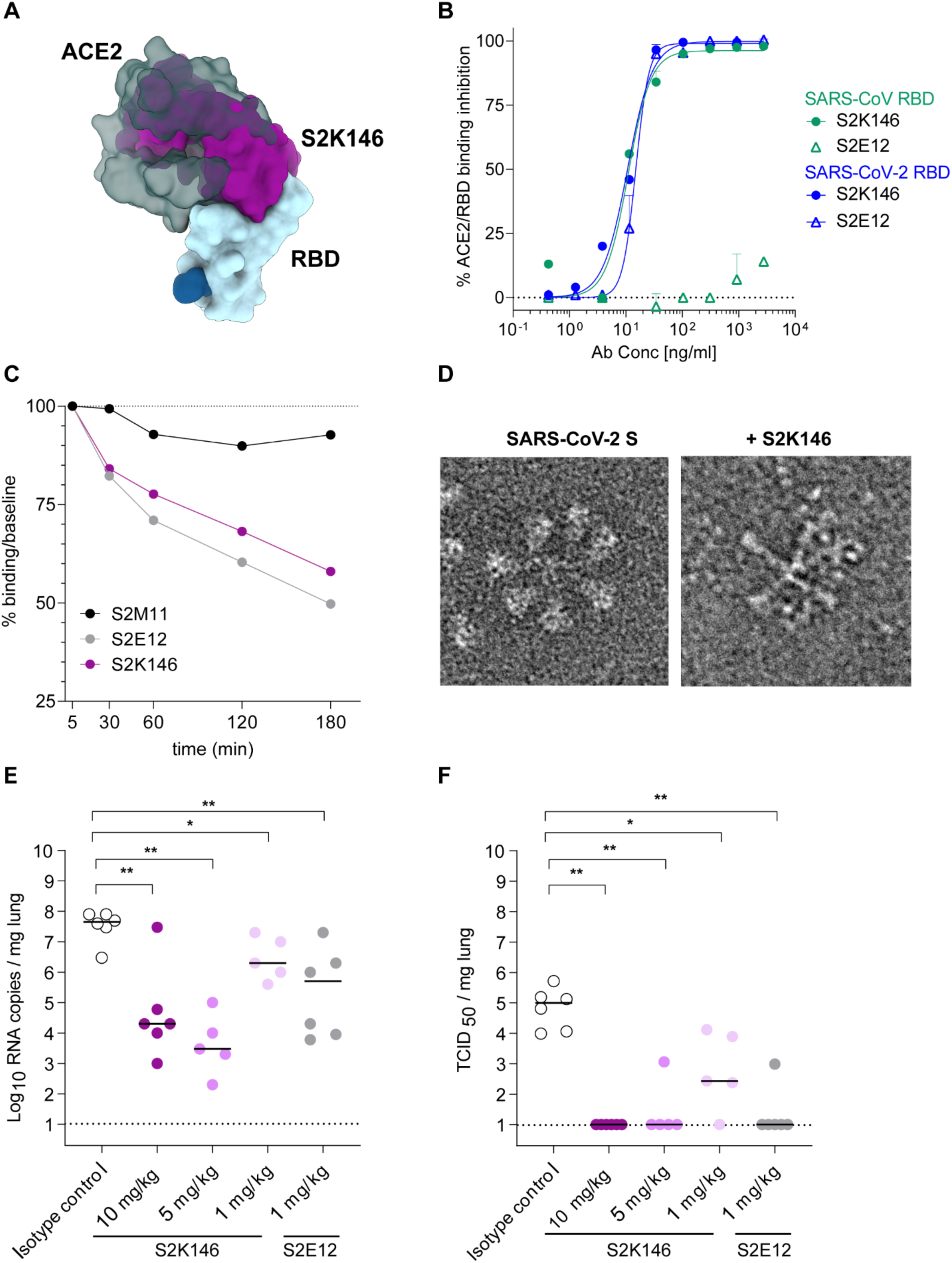
S2K146 blocks receptor attachment, triggers premature S refolding and protects against SARS-CoV-2 challenge therapeutically. **A)** Superimposition of the S2K146-bound (purple) and ACE2-bound (green, PDB: 6M0J (*36*)) SARS-CoV-2 RBD structures showing steric overlap. **B)** Pre-incubation of serial dilutions of S2K146 with SARS-CoV-2 RBD prevents binding to immobilized human ACE2 (hACE2) ectodomain in ELISA. **C)** S2K146-mediated S_1_-shedding from cell-surface expressed SARS-CoV-2 S as determined by flow cytometry. S2E12 mAb was used as positive control whereas S2M11 was used as a negative control. **D)** Cropped electron micrographs of negatively stained SARS-CoV-2 S trimer before (left, prefusion state) or after (right, postfusion state) incubation with S2K146. One representative micrograph for each dataset is shown out of 93 (SARS-CoV-2 S alone) and of 225 (SARS-CoV-2 S with S2K146) micrographs. **E-F)** Quantification of viral RNA (E) and replicating virus titers (TCID50), (F) in the lung of Syrian hamster 4 days post-intranasal infection with SARS-CoV-2 Beta VOC following therapeutic administration of S2K146 mAb at three different doses: 10-5-1mg/Kg (n=6/5 animal for each group). S2E12 mAb was administrated as control (n=6 animals). Isotype control was administered at 10 mg/kg (n=6 animals).

The efficient S2K146-induced S_1_ shedding could explain the lack of FcγRIIa and FcγRIIIa activation, which we used as a proxy for Ab-dependent cellular phagocytosis and Ab-dependent cellular cytotoxicity, respectively **(Fig. S7A-B)**. However, even when performing the same assays using target cells expressing an uncleavable prefusion stabilized SARS-CoV-2 S protein (unable to release the S_1_ subunit), S2K146 was not able to activate FcγRIIa and triggered FcγRIIIa weakly **(Fig. S7C-D)**. The greater efficiency of S2E12 for activating FcγRIIIa, relative to S2K146, might be explained by the different angles of approach of these two mAbs for binding to the RBD **(Fig. S7E-F)**.

Next, we evaluated the therapeutic activity of S2K146 against challenge with the Beta SARS-CoV-2 VOC in a Syrian hamster model of infection (*34, 35*). S2K146 was administered at 1, 5 and 10 mg/kg via intraperitoneal injection 24h after intranasal challenge with SARS-CoV-2 and the lungs of the animals were collected 3 days later for the quantification of viral RNA and replicating virus. In parallel, 6 animals were administered 1 mg/kg of the best-in-class, ultrapotent S2E12 mAb for benchmarking (*27*). Viral RNA loads in the lungs were reduced more than 1.5, 3 and 4 orders of magnitude after receiving 1, 5 and 10 mg/kg of S2K146, respectively **(Fig 4E)**. Viral replication in the lungs was completely abrogated for the 5 and 10 mg/kg groups and reduced by greater than 2.5 orders of magnitude for the 1 mg/kg group **(Fig 4F)**. Overall serum mAb concentrations measured at day 4 post-infection inversely correlated with viral RNA loads and infectious virus in the lungs **(Fig S8A-B)**. S2K146 therefore effectively protects against SARS-CoV-2 challenge in vivo in a stringent therapeutic setting.

## Discussion

The SARS-CoV-2 RBD accounts for most serum neutralizing activity in both COVID-19 convalescent (*11, 37*) and vaccinated individuals (*12*) and comprises antigenic sites targeted by broadly neutralizing sarbecovirus Abs (*13–19*). A few RBD-based subunit vaccines and mRNA vaccines based on chimeric S glycoproteins were recently shown to elicit broadly neutralizing sarbecovirus Abs and heterotypic protection in vivo (*38–42*). Most of these Abs with broad neutralizing activity are expected to target conserved RBD epitopes, due to their much greater potency and protection efficacy compared to known fusion machinery-directed Abs (*43–47*). The discovery of a functionally constrained and conserved RBM epitope associated with broad sarbecovirus neutralization rationalizes the strong cross-reactivity with the SARS-CoV RBM observed with polyclonal Abs elicited by a clinical stage SARS-CoV-2 vaccine in non-human primates (*40*) and will guide the development of next-generation pan-sarbecovirus vaccines to protect from future zoonotic transmission events.

The broadly neutralizing sarbecovirus mAb S309 was isolated from a survivor of a 2003 SARS-CoV infection and its derivative (sotrovimab) has received emergency use authorizations in several countries around the world for the early treatment of mild-to moderate COVID-19 in adults and pediatric patients (12 years of age and older weighing at least 40 kg) with positive results of direct SARS-CoV-2 viral testing, and who are at high risk for progression to severe COVID-19, including hospitalization or death (*13, 21, 23, 48*). S309 mAb has to date proven resilient to the emergence of SARS-CoV-2 variants, in pre-clinical studies, possibly due to targeting of a conserved RBD epitope with very limited mutational tolerance, potentially explaining its clinical success (*14*). The unique mechanism of S2K146-mediated ACE2 molecular mimicry provides an even higher barrier than S309 for emergence of escape mutants in spite of the known mutational plasticity of the SARS-CoV-2 RBM (*24*). Therefore, the discovery of the S2K146 mAb may be a key milestone for treatment of COVID-19 patients and for pandemic preparedness against divergent sarbecoviruses.

## Methods

### Cell lines

Cell lines used in this study were obtained from ATCC (Vero-E6) or ThermoFisher Scientific (Expi CHO cells and Expi293F™ cells) or were generated via lentiviral transduction (Expi CHO-S, HEK293T-ACE2, Vero-TMPRSS2) (*32*).

### Recombinant protein production

RBDs from different sarbecoviruses used in ELISA and BLI experiments were expressed with N-terminal signal peptide and C-terminal thrombin cleavage siteTwinStrep-8xHis-tag in Expi293F cells at 37°C and 8% CO_2_. Cells were transfected using PEI MAX (Polysciences) at a DNA:PEI ratio of 1:3.75. Transfected cells were supplemented three days after transfection with 3 g/L glucose (Bioconcept) and 5 g/L soy hydrolysate (Sigma-Aldrich Chemie GmbH). Cell culture supernatant (423 mL) was collected seven days after transfection and supplemented with 47 mL 10x binding buffer (1 M Tris-HCl, 1.5 M NaCl, 20 mM EDTA, pH 8.0) and 25 mL BioLock (IBA GmbH) and incubated on ice for 30 min. Proteins were purified using a 5 mL Strep-Tactin XT Superflow high capacity cartridge (IBA GmbH) followed by buffer exchange to PBS using HiPrep 26/10 desalting columns (Cytiva).

SARS-CoV-2 S hexapro (*49*), used for cryo-EM single particle studies, was expressed and purified as described before (*27*).

The SARS-CoV-2 S ‘wildtype’ ectodomain trimer used for refolding experiments followed by negative stain EM was engineered as follows and recombinantly expressed as previously described (*32*). The SARS-CoV-2 S D614G ‘wildtype’ has a mu-phosphatase signal peptide ending in ETGT, begins at Q14, a mutated S1/S2 cleavage site (SGAR), ends at residue K1211 and is followed by a TEV cleavage site, fold-on trimerization motif, and an 8× His tag in the pCMV vector.

### Antibody isolation and recombinant production

Antigen specific IgG^+^ memory B cells were isolated and cloned from PBMC of SARS-CoV-2 convalescent individuals. Briefly, CD19^+^ B cells were enriched by staining with CD19 PE-Cy7 and anti-PE microbeads (Milteniy), followed by positive selection using LS columns. Enriched B cells were stained with anti-IgD, anti-IgM, anti-IgA, anti-CD14, all PE labelled and prefusion SARS-CoV-2 S-Avi tag conjugated with streptavidin Alexa-Fluor 647 (Life Technologies). SARS-CoV-2-specific IgG^+^ memory B cells were sorted and seeded on MSC (mesenchymal stromal cells) at 0.5 cell/well in the presence of CpG2006, IL-2, IL6, IL-10 and IL-21, as previously described (*50*). After 7 days, B cell supernatants were screened by ELISA for binding to a panel of RBDs representative of different sarbecovirus clades as well as by neutralization using high-throughput VSV SARS-CoV-2 S-abesed microneutralization. Abs VH and VL sequences were obtained by reverse transcription PCR (RT-PCR) and mAbs were expressed as recombinant human IgG1, carrying the half-life extending M428L/N434S (LS) mutation in the Fc region fragment. ExpiCHO cells were transiently transfected with heavy and light chain expression vectors as previously described (*13*). Using the Database IMGT (http://www.imgt.org), the VH and VL gene family and the number of somatic mutations were determined by analyzing the homology of the VH and VL sequences to known human V, D and J genes. UCA sequences of heavy and light variable regions were constructed using IMGT/V-QUEST.

MAbs affinity purification was performed on ÄKTA Xpress FPLC (Cytiva) operated by UNICORN software version 5.11 (Build 407) using HiTrap Protein A columns (Cytiva) for full length human and hamster mAbs and CaptureSelect CH1-XL MiniChrom columns (ThermoFisher Scientific) for Fab fragments, using PBS as mobile phase. Buffer exchange to the appropriate formulation buffer was performed with a HiTrap Fast desalting column (Cytiva). The final products were sterilized by filtration through 0.22 μm filters and stored at 4ºC

### Enzyme-linked immunosorbent assay

96 half area well-plates (Corning, 3690) were coated over-night at 4°C with 25 μl of sarbecoviruses RBD proteins WIV1 (AGZ48831.1), Anlong-112 (ARI44804.1), YN2013 (AIA62330.1), SC2018 (QDF43815.1), ZC45 (AVP78031.1), Rp/Shaanxi2011 (AGC74165.1), BM48-31/BGR/2008 (YP_003858584.1), RaTG13 (QHR63300.2), SARS-CoV2 (YP_009724390.1), SARS-CoV Urbani (AAP13441.1), BtKY72 (APO40579.1), Pangolin-Guangdong-2019 (EPI_ISL_410721), Pangolin_Guanxi-2017 (EPI_ISL_410539) and SARS-CoV-2 RBD mutants, prepared at 5 μg/ml in PBS pH 7.2. After a blocking step of 60 min at room temperature with PBS 1% BSA (Sigma-Aldrich, A3059), plates were incubated with mAb serial dilutions for 60 min at room temperature. After 4 washing steps with PBS 0.05% Tween 20 (PBS-T) (Sigma-Aldrich, 93773), goat anti-human IgG secondary antibody (Southern Biotech, 2040-04) was added and incubated for 45 min at room temperature. Plates were then washed 4 times with PBS-T and 4-NitroPhenyl phosphate (pNPP, Sigma-Aldrich, 71768) substrate was added. After 45 min incubation, absorbance at 405 nm was measured by a plate reader (Biotek) and data plotted using Prism GraphPad.

### Transient Expression of sarbecovirus S protein in ExpiCHO-S Cells

ExpiCHO cells were seeded at 6 × 10^6^ cells/ml into 50 ml bioreactor tubes in 5 ml culture medium. Spike coding plasmids (5 μg) were diluted in OptiPRO SFM, mixed with ExpiFectamine CHO Reagent (Life Technologies) and added to the cells. After transfection, cells were incubated at 37°C with 8% CO_2_ with an orbital shaking speed of 120 rpm (orbital diameter of 25 mm) for 48 hours.

### Binding to cell surface expressed sarbecovirus S proteins by flow cytometry

Transiently transfected ExpiCHO cells were harvested and washed in wash buffer (PBS 2% FBS, 2 mM EDTA). Cells were counted, distributed into round bottom 96-well plates (Corning) and incubated with serially diluted antibodies in wash buffer (starting concentration: 10 μg/ml, 8 points of dilution 1:4). Alexa Fluor647-labeled Goat Anti-Human IgG secondary Ab (Jackson Immunoresearch) was prepared at 2 μg/mL added onto cells after two washing steps. Cells were then washed twice and resuspended in wash buffer for data acquisition at ZE5 cytometer (Biorad).

### Competition assay and affinity determination by Biolayer Interferometry (BLI)

BLI experiments were carried out using an Octet Red96 (ForteBio) and all reagents were prepared in Kinetics buffer (KB) (PBS 0.01% BSA).

To assess S2K146 competition with S2×259, S309 and S2E12, His-tagged SARS-CoV-2 RBD was prepared at 8 μg/ml in Kinetics buffer (PBS 0.01% BSA) and loaded on pre-hydrated anti-penta-HIS biosensors (Sartorius) for 2.5 min. Biosensors were then moved into a solution containing 20 μg/ml S2K146 mAb and association recorded for 5 min. A second association step was subsequently performed into S2×259, S309 and S2E12 mAbs solutions at 20 μg/ml and recorded for 5 min. Response values were exported and plotted using GraphPad Prism (version 9.1.1).

To assess binding affinities, S2K146 and respective UCA Ab were prepared at 3 μg/ml and immobilized on pre-hydrated protein-A biosensors (Sartorius) for 75 sec. After a 30 sec stabilization step in KB, biosensors were moved in SARS-CoV or SARS-CoV-2 :2 dilution series (starting concentration: 18.5 nM) for the 600 sec association step, and then moved back in KB to record dissociation signals for 540 sec. The data were baseline subtracted, results fitted using the Pall FortéBio/Sartorius analysis software (version 12.0) and plotted using GraphPad Prism (version 9.1.1)

### VSV-based pseudovirus production and neutralization assay

VSV psudoviruses were produced using the following constructs: SARS-CoV S, SARS-CoV-2 S, WIV-1 S, RaTG13 S, PG-GD S, PG-GX S, the VOC Alfa, Beta, Gamma, Epsilon, the VOI B.1.1.519, B.1.1.218 and SARS-CoV2 S bearing the single mutations K417V, E484K. Pseudotyped viruses were prepared using Lenti-X 293 cells seeded in 15-cm dishes. Briefly, cells in culture medium (DMEM supplemented with 10% heat-inactivated FBS, 1% PenStrep) were transfected with 25 μg of plasmid encoding for the corresponding S glycoprotein using TransIT-Lenti (Mirus) as transfectant reagent. One day post-transfection, cells were infected with VSV (G*ΔG-luciferase) for 1 h, washed 3 times in PBS with Ca^2+^/Mg^2+^ (Thermo Fisher) before adding 25 ml of culture medium/dish. Particles were harvested after 18-24 h, clarified from cellular debris by centrifugation at 2,000 × g for 20 min at 4°C, aliquoted and stored at −80°C until use in neutralization experiments.

For neutralization experiments, Vero E6 cells were seeded at 20,000 cells/well in culture medium into white 96-well plates (PerkinElmer, 6005688) and cultured overnight at 37°C 5% CO_2_. Ten-point 3-fold mAb serial dilutions were prepared in culture medium and mixed 1:1 with pseudotyped VSV prepared in culture medium in order to infect cells with the desired MOI. After 60 min incubation at 37 °C, cell culture medium was aspirated and 50 μl of PVs/mAb mixture was added onto cells and incubated 60 min at 37°C 5% CO_2_. After 60 min, 100 μl of culture medium was added to the cells and incubation at 37°C 5% CO_2_ followed in the next 16-24 h. At the end of the incubation time, culture medium was removed from the cells and 50 μl/well of Steadylite (PerkinElmer) diluted 1:2 with PBS with Ca^2+^Mg^2+^ was added to the cells and incubated in the dark for 10 min. Luminescence signals were read using a Synergy H1 Hybrid Multi-Mode plate reader (Biotek). Measurements were done in duplicate and at least six wells per plate contained untreated infected cells (defining the 0% of neutralization, “MAX RLU” value) and infected cells in the presence of S2E12 and S2×259 at 25 μg/ml each (defining the 100% of neutralization, “MIN RLU” value). Average of Relative light units (RLUs) of untreated infected wells (MAX RLUave) was subtracted by the average of MIN RLU (MIN RLUave) and used to normalize percentage of neutralization of individual RLU values of experimental data according to the following formula: (1-(RLUx - MIN RLUave) / (MAX RLUave – MIN RLUave)) × 100. Data were analyzed and visualized with Prism (Version 9.1.1). IC50 values were calculated from the interpolated value from the log(inhibitor) versus response, using variable slope (four parameters) nonlinear regression with an upper constraint of ≤100, and a lower constrain equal to 0.

### BtKY72 (K493Y/T498W) S pseudovirus production and neutralization assay

VSV pseudovirus harboring BtKY72 K493Y/T498W S (mutants defined based on SARS-CoV-2 numbering) with a native signal peptide and C-terminal 21 residue deletion synthesized by GenScript were prepared as previously described (*28*). Briefly, HEK-293T cells seeded in poly-D-lysine coated 100 mm dishes at ~75 % confluency were washed five times with Opti-MEM and co-transfected with Lipofectamine 2000 (Life Technologies) with 24 μg of the S glycoprotein plasmids. After 5 h at 37°C, media supplemented with 20% FBS and 2% PenStrep was added. After 20 hours, cells were washed five times with DMEM and cells were transduced with VSVΔG-luc (*51*) and incubated at 37°C. After 2 h, infected cells were washed an additional five times with DMEM prior to adding media supplemented with anti-VSV-G antibody (I1-mouse hybridoma supernatant diluted 1:25, from CRL-2700, ATCC) to reduce parental background. After 18-24 h, the supernatant was harvested and clarified by low-speed centrifugation at 2,500 g for 10 min. The supernatant was then filtered (0.45 μm) and concentrated 10 times using a 30 kDa cut off membrane. The pseudotypes were then aliquoted and frozen at −80 °C.

For neutralization experiments, HEK-293T cells expressing hACE2 (Crawford et al. 2020) in DMEM supplemented with 10% FBS and 1% PenStrep were seeded at 20,000 cells per well into clear bottom, white manually poly-D-lysine coated 96 well plates and incubated at 37°C. The following day, an additional half-area, 96-well plate was prepared with twelve 3-fold serial dilutions of mAb of either S2K146 or S2×259 starting at 250ug/mL or 25 ug/mL respectively. An equal volume of diluted pseudovirus was added and incubated for 30 minutes at room temperature. Excess media was removed from cells and the mAb-pseudovirus mixture was transferred to the cells for 2 hours at 37C. After the 2 hour incubation, an equal volume of DMEM-20%FBS-2%PenStrep was added for overnight incubation. The following day, One-Glo-EX substrate (Promega) was added and incubated in the dark for 5 minutes. The plates were immediately read on a Biotek plate reader. Relative luciferase units were plotted and normalized in Prism with cells alone without pseudovirus defining 100% neutralization and cells with pseudovirus only defining 0% neutralization. Data were analyzed and visualized with Prism (Version 9.1.1). IC50 values were calculated from the interpolated value from the log(inhibitor) versus response, using variable slope (four parameters) nonlinear regression with an upper constraint of ≤100, and a lower constrain equal to 0.

### Authentic SARS-CoV-2 isolates

SARS-CoV-2 strains used in this study were obtained from BEI (WT: Lineage A, BEI ref. NR-5228; Alpha: Lineage B.1.1.7, BEI ref. NR-54000; Beta: Lineage B.1.351, BEI ref. NR-54009; Kappa: Lineage B.1.617.1, BEI ref. NR-55486; Delta: Lineage B.1.617.2, BEI ref. NR-55611).

### Neutralization of authentic SARS-CoV-2 viruses

Vero-TMPRSS2 cells were seeded into black-walled, clear-bottom 96-well plates at 2 × 10^4^ cells/well and cultured overnight at 37°C. The next day, 9-point 5-fold serial dilutions of mAbs were prepared in infection media (DMEM + 10% FBS). The different SARS-CoV-2 strains were diluted in infection media at a final MOI of 0.01 PFU/cell, added to the mAb dilutions and incubated for 30 min at 37°C. Media was removed from the cells, mAb-virus complexes were added and incubated at 37°C for 18 h. Cells were fixed with 4% PFA (Electron Microscopy Sciences, #15714S), permeabilized with Triton X-100 (SIGMA, #X100-500ML) and stained with an antibody against the viral nucleocapsid protein (Sino Biologicals, #40143-R001) followed by a staining with the nuclear dye Hoechst 33342 (Fisher Scientific, # H1399) and a goat anti-rabbit Alexa Fluor 647 antibody (Invitrogen, #A-21245). Cells were imaged with an automated multimode plate reader (Biotek, Cytation 5).

### Blockade of SARS-CoV and SARS-CoV-2 binding to ACE2

SARS-CoV and SARS-CoV-2 mouse/rabbit Fc-tagged RBDs (final concentration 20 ng/ml) were incubated with serially diluted recombinant mAbs (from 25 μg/ml) and incubated for 1 h 37°C. The complex RBD:mAbs was then added to a pre-coated hACE2 (2 μg/ml in PBS) 96-well plate MaxiSorp (Nunc) and incubated 1 hour at room temperature. Subsequently, the plates were washed and a goat anti-mouse/rabbit IgG (Southern Biotech) coupled to alkaline phosphatase (Jackson Immunoresearch) added to detect mouse Fc-tagged RBDs binding. After further washing, the substrate (p-NPP, Sigma) was added, and plates read at 405 nm using a microplate reader (Biotek). The percentage of inhibition was calculated as follow: (1-((OD sample-OD neg ctr)/(OD pos. ctr-OD neg. ctr))*100

### Cell-surface mAb-mediated S_1_ shedding

CHO cells stably expressing the prototypic SARS-CoV-2 Spike protein were harvested, washed in wash buffer (PBS 1% BSA 2 mM EDTA) and resuspended in PBS. Cells were then counted and 90’000 cells/well were dispensed into a round-bottom 96 well plate (Corning) to be treated with 10 ug/ml TPCK-Trypsin (Worthington Biochem) for 30 min at 37°C. After a washing step, cells were incubated with 15 ug/ml mAbs solution for 180, 120, 60, 30 or 5 min at 37°C. After the incubation for the allotted time, cells were washed with ice-cold wash buffer and stained with 1.5 ug/ml Alexa Fluor647-labeled Goat Anti-Human IgG secondary Ab (Jackson Immunoresearch) for 30 min on ice in the dark. Cells were then washed twice with cold wash buffer and analyzed using a ZE5 cytometer (Biorad) with acquisition chamber T= 4°C. Binding at each time point (MFI) was determined normalizing to the MFI at 5 minutes time point and data plotted using GraphPad Prism v. 9.1.1

### Evaluation of escape mutants via deep mutational scanning

A previously described deep mutational scanning approach (*52*) was used to identify RBD mutations that escape S2K146 binding exactly as described in (*14*). Briefly, duplicate libraries containing virtually all possible amino acid changes compatible with ACE2 binding and RBD folding within the Wuhan-Hu-1 SARS-CoV-2 RBD sequence were expressed on the surface of yeast. Libraries were labelled at 63 ng/mL S2K146 antibody (the EC90 for binding to yeast-displayed SARS-CoV-2 RBD determined in isogenic pilot binding experiments), and fluorescence-activated cell sorting (FACS) was used to select RBD+ cells that exhibit reduced antibody binding as previously described. Libraries were sequenced before and after selection to determine per-mutation escape fractions as previously described. Experiments were performed in duplicate with independently generated mutant libraries, and we report the average mutant escape fraction across the duplicates. Data for S2E12 exactly as described in (*14*) are included for comparison to S2K146. Complete computational pipeline for deep mutational scanning data analysis is available on GitHub: https://github.com/jbloomlab/SARS-CoV-2-RBD_MAP_S2K146.

### Evaluation of sarbecovirus cross-reactivity via high-throughput yeast-display binding assays

Breadth of antibody binding across a panel of yeast-displayed sarbecovirus RBDs was performed as described in (cites: PMID 34261126), with modification. In the prior publication, antibody binding to the sarbecovirus RBD panel was measured at a single antibody concentration analogous to the deep mutational scanning selections. In the current work, binding was determined via FACS-seq across an antibody dilution series (10,000 to 0.01 ng/mL in 10-fold dilutions, plus 0 ng/mL antibody). The ‘escape fraction’ of each sarbecovirus RBD at each antibody concentration was fit to a sigmoid binding curve to determine the quantitative EC50 for antibody binding to each RBD. In addition to S2K146 breadth, S2E12 was profiled using this new method in this paper to enable direct comparison to S2K146. Complete computational pipeline for the analysis of breadth of antibody binding is available on GitHub: https://github.com/jbloomlab/SARSr-CoV_RBD_MAP. Several sarbecovirus RBD sequences were reported subsequent to the cloning of our original sarbecovirus panel, including RshSTT182 (cites: doi: 10.1101/2021.01.26.428212, GISAID EPI_ISL_852604), Rc-o319 (cites: PMID 33219796, Genbank: LC556375), PRD-0038 and PDF-2370 (cite:, Genbank MT726045 and MY726044), and RsYN04 (cite: PMID 34147139, GISAID EPI_ISL_1699444). These RBD sequences were cloned into the yeast-surface display vector and antibody binding was determined via isogenic binding assays monitored by flow cytometry, exactly as described in (*14*).

### Selection of SARS-CoV-2 monoclonal antibody escape mutants

A VSV-SARS-CoV-2 Wuhan-Hu-1 D614G S chimera was used to select for mAb resistant mutants, as previously described(*30*). Briefly, mutants were recovered by plaque isolation on Vero E6 cells with the indicated mAb in the overlay. The concentration of mAb in the overlay was determined by neutralization assays at a multiplicity of infection (MOI) of 100. Escape clones were plaque-purified on Vero E6 cells in the presence of mAb, and plaques in agarose plugs were amplified on MA104 cells with the mAb present in the medium. Viral stocks were amplified on MA104 cells at an MOI of 0.01 in Medium 199 containing 2% FBS and 20 mM HEPES pH 7.7 (Millipore Sigma) at 34°C. Viral supernatants were harvested upon extensive cytopathic effect and clarified of cell debris by centrifugation at 1,000 × g for 5 min. Aliquots were maintained at −80°C. Viral RNA was extracted from VSV-SARS-CoV-2 S mutant viruses using RNeasy Mini kit (Qiagen), and the S gene was amplified using OneStep RT-PCR Kit (Qiagen). The mutations were identified by Sanger sequencing (GENEWIZ). Their resistance was verified by subsequent virus infection in the presence or absence of mAb. Vero E6 cells were seeded into 12 well plates overnight. The virus was serially diluted using DMEM and cells were infected at 37°C for 1 h. Cells were cultured with an agarose overlay in the presence or absence of mAb at 34°C for 2 days. Plates were scanned on a biomolecular imager and expression of eGFP monitored at 48 hours post-infection.

### Viral replication fitness assays

Vero E6 cells (ATCC, CRL-1586) were seeded at 1×10^6^ cells per well in 6-well plates. Cells were infected with multiplicity of infection (MOI) of 0.02, with VSV chimeras harboring SARS-CoV-2 Wuhan-Hu-1 D614G S and SARS-CoV-2 Wuhan-Hu-1 Y489H/D614G S mixed at equal titers. Following 1 h incubation, cell monolayers were washed three times with Hanks’ Balanced Salt Solution (HBBS) and cultures were incubated for 72 h in humidified incubators at 34°C. To passage the progeny viruses, virus mixture was continuously passaged four times in Vero E6 cells at MOI of 0.02. Cellular RNA samples from each passage were extracted using RNeasy Mini kit (QIAGEN) and subjected to next-generation sequencing as described previously to confirm the introduction and frequency of substitutions (*30*).

### S2K146-induced S refolding

10 μM native-like SARS-CoV-2 S(*32*) was incubated with 13 μM S2K146 Fab for 12 hours at room temperature. Samples were diluted to 0.01 mg/mL immediately prior to adsorption to glow-discharged carbon-coated copper grids for ~30 sec prior to a 2% uranyl formate staining. Micrographs were recorded using the Leginon software on a 120 kV FEI Tecnai G2 Spirit with a Gatan Ultrascan 4000 4k × 4k CCD camera at 67,000 nominal magnification. The defocus ranged from −1.0 to −2.0 μm and the pixel size was 1.6 Å.

### CryoEM sample preparation, data collection and data processing

Recombinantly expressed and purified S2K146 Fab and SARS-CoV-2 S hexapro(*49*) were incubated at 1 mg/ml (for UltraAuFoil grids) or 0.1mg/ml (for lacey thin carbon grids) with a 1.2 molar excess of Fab at 4°C for 1 hr. Three microliters of the complex mixture were loaded onto freshly glow discharged R 2/2 UltrAuFoil grids (200 mesh) or lacey grids covered with a thin layer of manually added carbon, prior to plunge freezing using a vitrobot MarkIV (ThermoFisher Scientific) with a blot force of 0 and 6-6.5 sec blot time (for the UltrAuFoil grids) or with a blot force of −1 and 2.5 sec blot time (for the lacey thin carbon grids) at 100 % humidity and 22°C.

Data were acquired using an FEI Titan Krios transmission electron microscope operated at 300 kV and equipped with a Gatan K3 direct detector and Gatan Quantum GIF energy filter, operated in zero-loss mode with a slit width of 20 eV. Automated data collection was carried out using Leginon (*53*) at a nominal magnification of 105,000x with a pixel size of 0.843 Å. The dose rate was adjusted to 15 counts/pixel/s, and each movie was acquired in super-resolution mode fractionated in 75 frames of 40 ms. 7,289 micrographs were collected with a defocus range comprised between −0.5 and −2.5 μm. Movie frame alignment, estimation of the microscope contrast-transfer function parameters, particle picking, and extraction were carried out using Warp (*54*).

Two rounds of reference-free 2D classification were performed using CryoSPARC (*55*) to select well-defined particle images. These selected particles were subjected to two rounds of 3D classification with 50 iterations each (angular sampling 7.5° for 25 iterations and 1.8° with local search for 25 iterations), using our previously reported closed SARS-CoV-2 S structure as initial model (PDB 6VXX) (*4*) using Relion (*56*). 3D refinements were carried out using non-uniform refinement along with per-particle defocus refinement in CryoSPARC (*57*). Selected particle images were subjected to the Bayesian polishing procedure (*58*) implemented in Relion3.0 before performing another round of non-uniform refinement in cryoSPARC followed by per-particle defocus refinement and again non-uniform refinement. To improve the density of the S/S2K146 interface, the particles from the class with 2 RBDs opened were subjected to focus 3D classification without refining angles and shifts using a soft mask on the closed RBD and bound S2K146 variable domains with a tau value of 60 in Relion. Particles belonging to classes with the best resolved local density were selected and subject to local refinement using CryoSPARC. Local resolution estimation, filtering, and sharpening were carried out using CryoSPARC. Reported resolutions are based on the gold-standard Fourier shell correlation (FSC) of 0.143 criterion and Fourier shell correlation curves were corrected for the effects of soft masking by high-resolution noise substitution (*59, 60*).

### Model building and refinement

UCSF Chimera (*61*) and Coot (*62*) were used to fit atomic models into the cryoEM maps. Spike-RBD/S2K146 model was refined and relaxed using Rosetta using sharpened and unsharpened maps (*63, 64*).

### Measurement of Fc-effector functions

S2K146-dependent activation of human FcγRIIa and IIIa was performed with a bioluminescent reporter assay. ExpiCHO cells stably expressing full-length wild-type SARS-CoV-2 S (target cells) or full-length prefusion stabilized SARS-CoV-2 S, which harbours the 2P mutation and S1/S2 furin cleavage site mutation (RRARS to SGAG) as previously described (*4*), were incubated with different amounts of mAbs. After a 15-minute incubation, Jurkat cells stably expressing FcγRIIIa receptor (V158 variant) or FcγRIIa receptor (H131 variant) and NFAT-driven luciferase gene (effector cells) were added at an effector to target ratio of 6:1 for FcγRIIIa and 5:1 for FcγRIIa. Signaling was quantified by the luciferase signal produced as a result of NFAT pathway activation. Luminescence was measured after 20 hours of incubation at 37°C with 5% CO_2_ with a luminometer using the Bio-Glo-TM Luciferase Assay Reagent according to the manufacturer’s instructions (Promega).

### Hamster challenge experiment

The hamster infection model of SARS-CoV-2 including the associated analytical procedures, have been described before (*35, 65*). In brief, female Syrian hamsters (Mesocricetus auratus) of 6-8 weeks old were anesthetized with ketamine/xylazine/atropine and inoculated intranasally with 50 μL containing 1×10^4^ TCID50 Beta B.1.351 (derived from hCoV-19/Belgium/rega-1920/2021; EPI_ISL_896474, 2021-01-11). This variant was originally isolated in house from nasopharyngeal swabs taken from travelers returning to Belgium (baseline surveillance) and were subjected to sequencing on a MinION platform (Oxford Nanopore) directly from the nasopharyngeal swabs (*65*). Animals were treated once by intraperitoneal injection 24h post SARS-CoV-2 challenge (i.e. therapeutic administration) with S2K146 mAb (at 10, 5 and 1 mg/Kg). S2E12 mAb was administrated as control (at 1 mg/kg). Isotype control was administered at 10 mg/kg. Hamsters were monitored for appearance, behavior and weight. At day 4 pi, hamsters were euthanized by i.p. injection of 500 μL Dolethal (200mg/ml sodium pentobarbital, Vétoquinol SA). Lungs were collected for viral RNA and infectious virus quantification by RT-qPCR and end-point virus titration, respectively. Serum samples were collected at day 4 pi for analysis of Ab levels.

### SARS-CoV-2 RT-qPCR

Hamster lung tissues were collected after sacrifice and were homogenized using bead disruption (Precellys) in 350 μL TRK lysis buffer (E.Z.N.A.^®^ Total RNA Kit, Omega Bio-tek) and centrifuged (10.000 rpm, 5 min) to pellet the cell debris. RNA was extracted according to the manufacturer’s instructions. RT-qPCR was performed on a LightCycler96 platform (Roche) using the iTaq Universal Probes One-Step RT-qPCR kit (BioRad) with N2 primers and probes targeting the nucleocapsid (*65*). Standards of SARS-CoV-2 cDNA (IDT) were used to express viral genome copies per mg tissue.

### End-point virus titrations

Lung tissues were homogenized using bead disruption (Precellys) in 350 μL minimal essential medium and centrifuged (10,000 rpm, 5min, 4°C) to pellet the cell debris. To quantify infectious SARS-CoV-2 particles, endpoint titrations were performed on confluent Vero E6 cells in 96-well plates. Viral titers were calculated by the Reed and Muench method (*66*) using the Lindenbach calculator and were expressed as 50% tissue culture infectious dose (TCID_50_) per mg tissue.

## ACKNOWLEDGMENTS

The authors thank Cindy Castado and Normand Blais (GSK Vaccines) for their help in the selection of the genetically divergent sarbecoviruses used in this study and Hideki Tani (University of Toyama) for providing the reagents necessary for preparing VSV pseudotyped viruses. This study was supported by the National Institute of Allergy and Infectious Diseases (DP1AI158186 and HHSN272201700059C to D.V.), a Pew Biomedical Scholars Award (D.V.), an Investigators in the Pathogenesis of Infectious Disease Awards from the Burroughs Wellcome Fund (D.V.), Fast Grants (D.V.), the University of Washington Arnold and Mabel Beckman cryoEM center and the National Institute of Health grant S10OD032290 (to D.V.).

## Author contributions

Y.J.P, A.D.M, T.N.S., Z.L., D.P., J.D.B., D.C., M.S.P and D.V. designed the experiments; A.D.M, D.P., A.C.W., K.S.S. F.Z., M.G., J.N. and F.A.L isolated mAb and performed binding, neutralization assays and biolayer interferometry measurements; A.D.M. and D.P. performed ACE2 binding inhibition and S_1_ shedding assays; B.G. evaluated effector functions; T.N.S. and J.D.B. performed deep-mutational scanning; Z.L. and S.P.J.W. performed mutant selection and fitness assays; R.A., S.-Y.C.F., F.B., and J.N., D.C. and M.S.P. performed hamster model experiments and data analysis; Y.J.P. carried out cryoEM specimen preparation, data collection and processing. Y.J.P. and D.V. built and refined the atomic models. S.K.Z., A.J. and J.E.B. purified recombinant glycoproteins. Y.J.P., A.D.M., T.N.S., Z.L., D.P., J.D.B., D.C., M.S.P and D.V. analyzed the data; Y.J.P., A.D.M, D.C., M.S.P and D.V. wrote the manuscript with input from all authors; F.A.L., F.B., G.S., J.N., S.P.J.W., H.W.V., J.D.B., D.C., M.S.P. and D.V. supervised the project.

## Data and materials availability

The cryoEM map and coordinates have been deposited to the Electron Microscopy Databank and Protein Data Bank with accession numbers XXX, YYY, ZZZ.

## Supplemental figures

**Figure S1.**
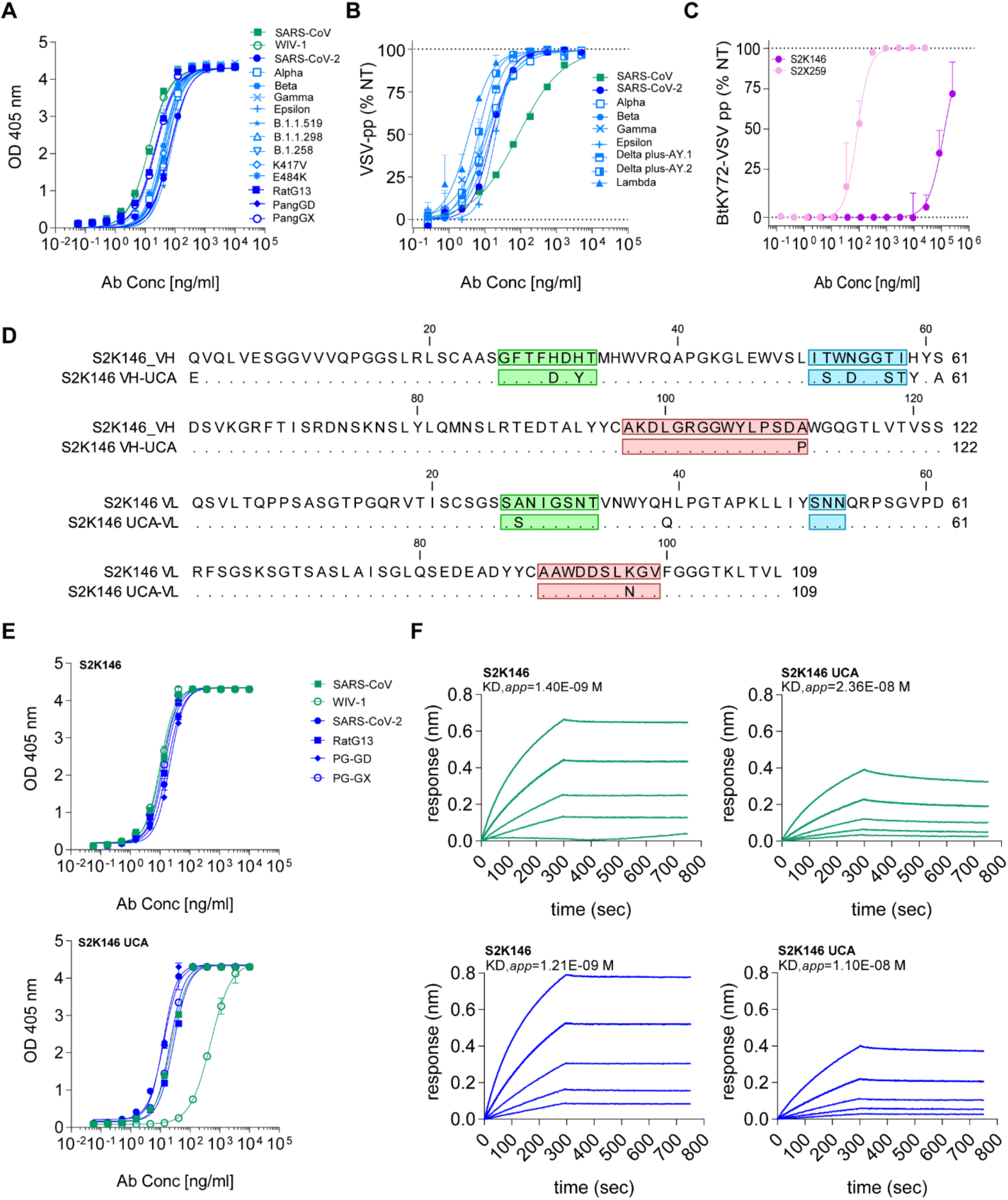
In vitro characterization of S2K146 mAb. **A)** ELISA binding curves of S2K146 mAb against different clade 1a and 1b sarbecovirus RBDs. Serial dilutions of S2K146 mAb were tested. **B)** S2K146-mediated neutralization of VSV pseudotypes harboring Wuhan-Hu-1 SARS-CoV-2 S, B.1.351 (beta), B.1.1.7 (alpha), P.1 (gamma), B.1.429 (epsilon), C.37 (lambda), AY.1/.2 (delta+) SARS-CoV-2 S or SARS-CoV S. **C)** S2K146-mediated neutralization of VSV pseudotypes harboring BtKY72 S (K493Y/T498W, SARS-CoV-2 residue numbering) using HEK293T cells expressing human ACE2. The S2×259 mAb was included as a positive control. **D**) Alignment of the amino acid sequence of the variable regions of heavy and light chains of S2K146 and S2K146 UCA. Heavy and light chain CDR1, CDR2, and CRD3 (IMGT definition) are indicated by green, blue, and red boxes, respectively. **E)** ELISA binding analysis of S2K146 and S2K146 UCA to RBDs of clade 1a and clade1b sarbecoviruses. **F)** Biolayer interferometry binding analysis of S2K146 and S2K146 UCA IgG1 to prefusion SARS-CoV S (green) and SARS-CoV-2 S (blue). Apparent KD (KD,*app*) are reported.

**Figure S2.**
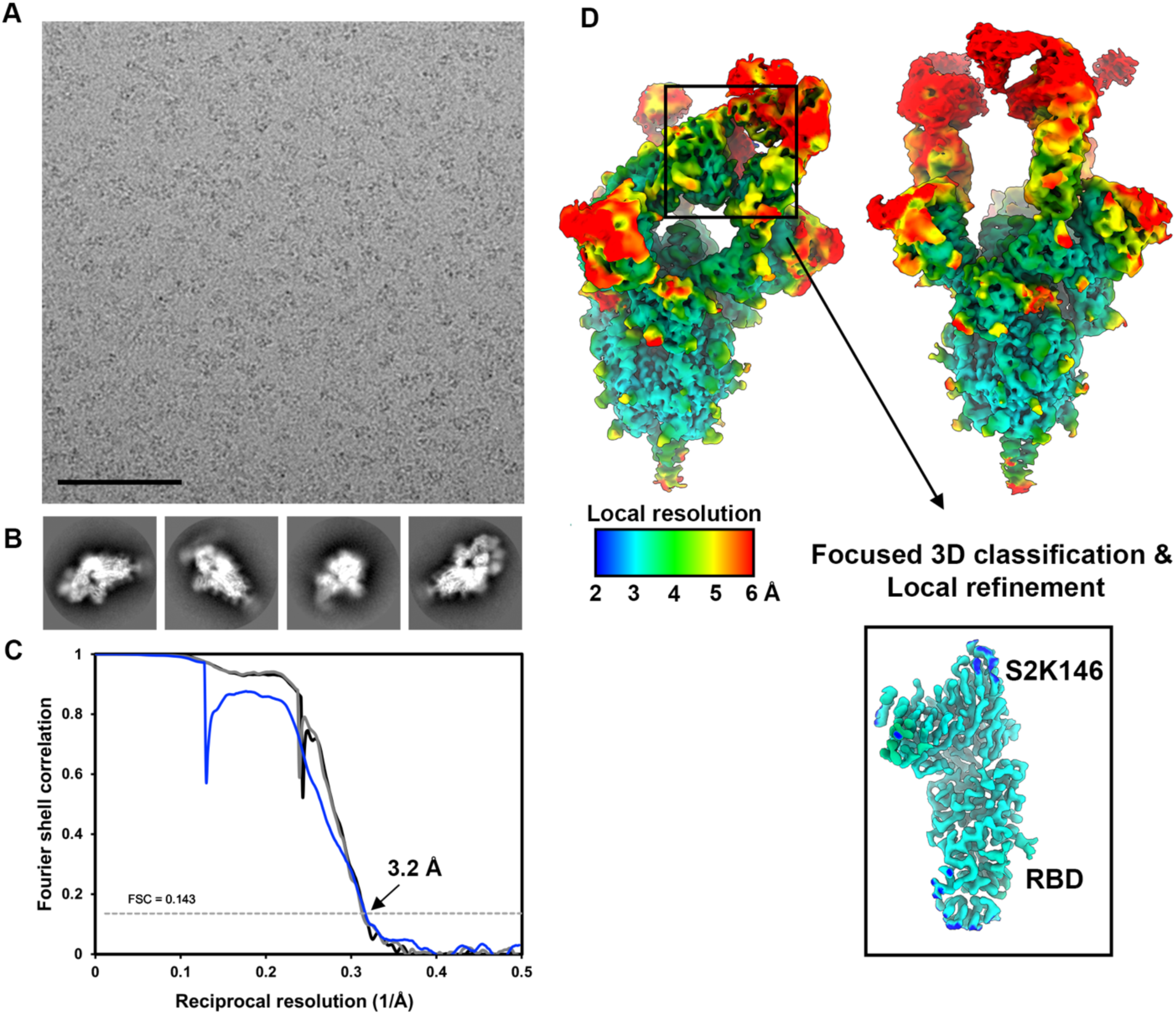
CryoEM data processing of the S2K146-bound SARS-CoV-2 S dataset. **A-B)** Representative electron micrograph and 2D class averages of SARS-CoV-2 S in complex with S2K146 Fab embedded in vitreous ice. The scale bar represents 100 nm. **C)** Gold-standard Fourier shell correlation curves for the S2K146-bound SARS-CoV-2 S maps with three RBDs open (black line), two RBDs open (grey line) and locally refined RBD/S2K146 variable domain (blue line). The 0.143 cutoff is indicated by a horizontal dashed line. **D)** Local resolution maps calculated using cryoSPARC for the whole reconstruction of two RBD in open state (left) and three RBD open state (right) SARS-CoV-2 S trimer as well as for the locally refined RBD/S2K146 variable domain region.

**Figure S3.**
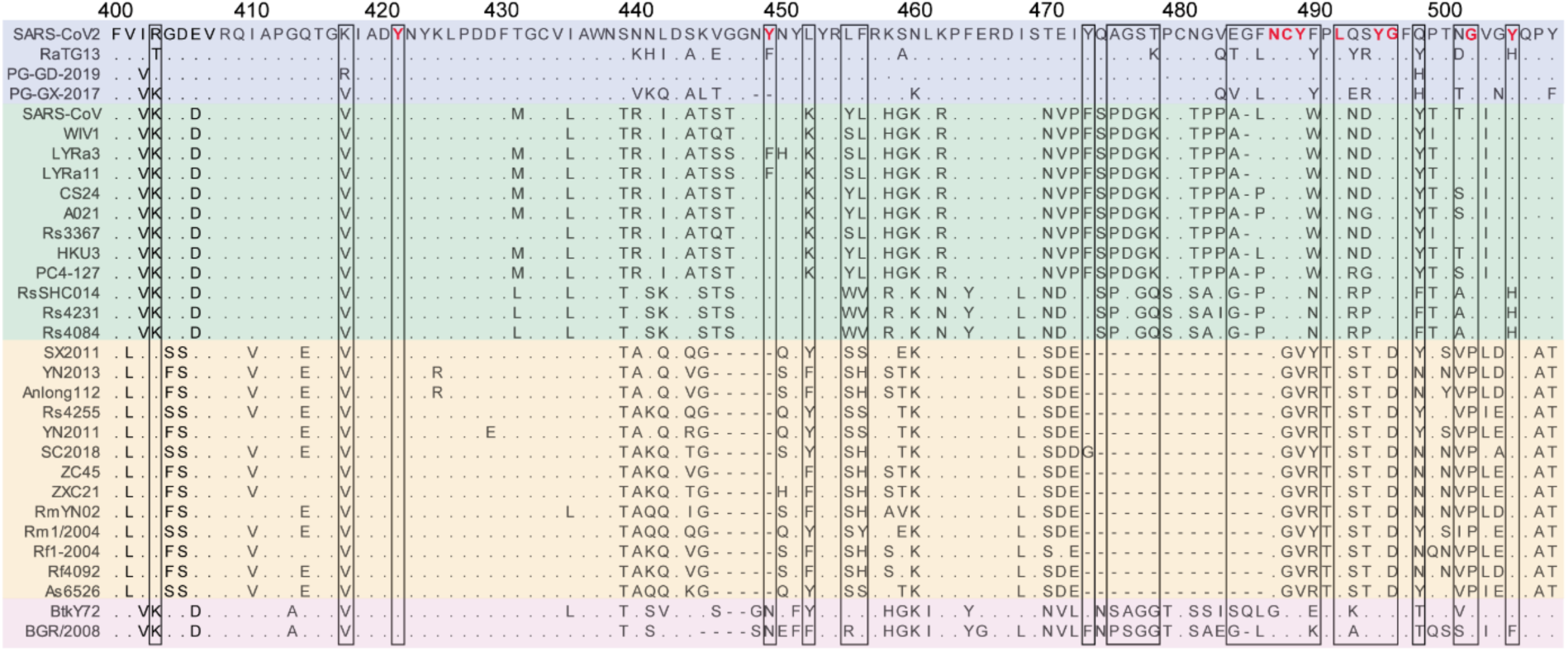
S2K146 epitope conservation across sarbecoviruses. Protein sequence alignment of RBDs representative of sarbecovirus clade 1a (green), clade 1b (blue), clade 2 (orange) and clade 3 (pink). Residue numbering is based on SARS-CoV-2 sequence with identical residues indicated as dots. Residues buried upon S2K146 binding are highlighted in boxes and epitope residues conserved between the SARS-CoV-2 and SARS-CoV RBDs are denoted in red.

**Figure S4.**
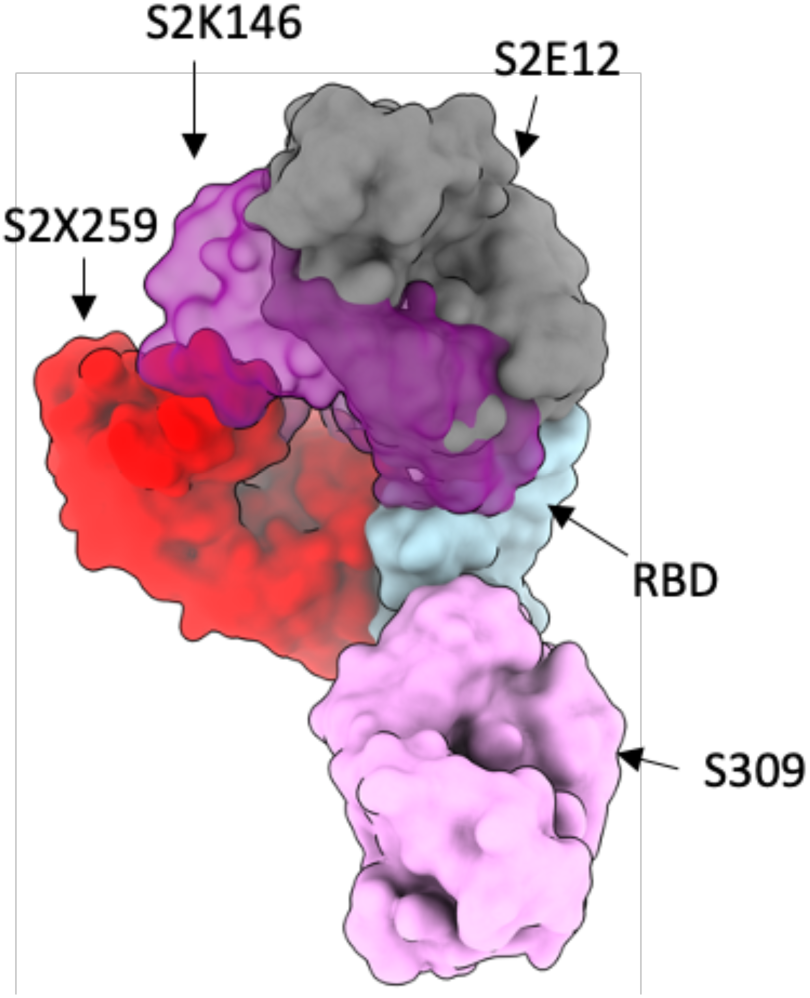
SARS-CoV-2 broadly neutralizing sarbecovirus mAbs. Structural superimposition of S2K146 (purple), site I-targeting S2E12 (grey), site II-targeting S2×259 (red), and site IV-targeting S309 (pink) Fabs bound to the SARS-CoV-2 RBD (light blue). S2K146 is depicted as a semi-transparent surface to show that S2E12 and S2K146 bind largely overlapping binding sites on the RBD. Although the constant domains of S2K146 and S2×259 slightly overlap, the flexibility around the Fab elbow allows binding of both mAbs as shown in Fig 1A.

**Figure S5.**
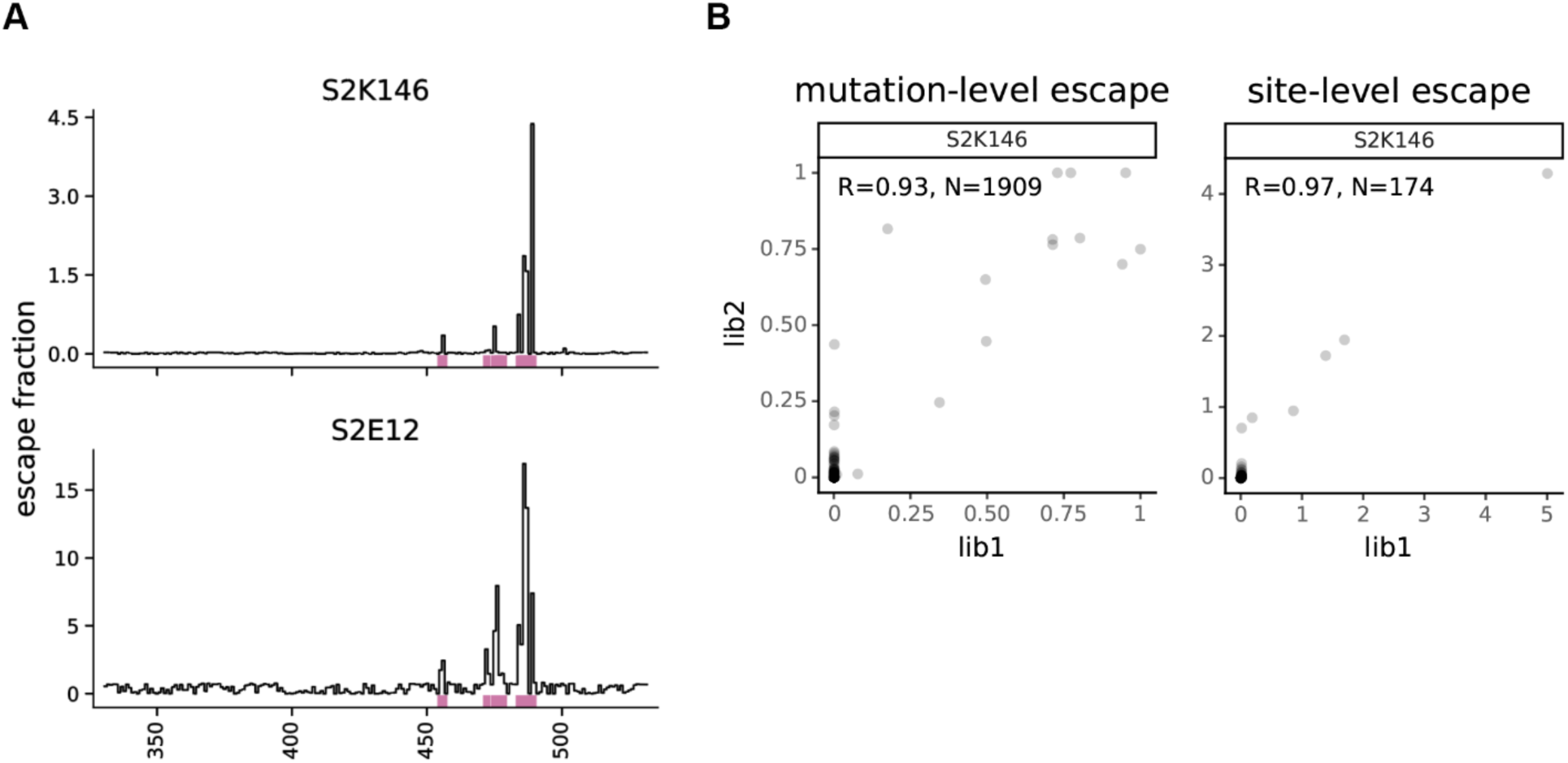
DMS analysis of the yeast-displayed SARS-CoV-2 RBD. **A)** Mapping of escape mutations reducing S2K146 or SE12 binding using yeast-displayed RBD DMS. Line plots show escape at each site in the RBD (summed effects of all mutations at each site). Sites of strong escape are indicated with purple lines. **B**) Correlation in per-mutation (left) and sum-per-site (right) escape fraction for replicate DMS library experiments.

**Figure S6.**
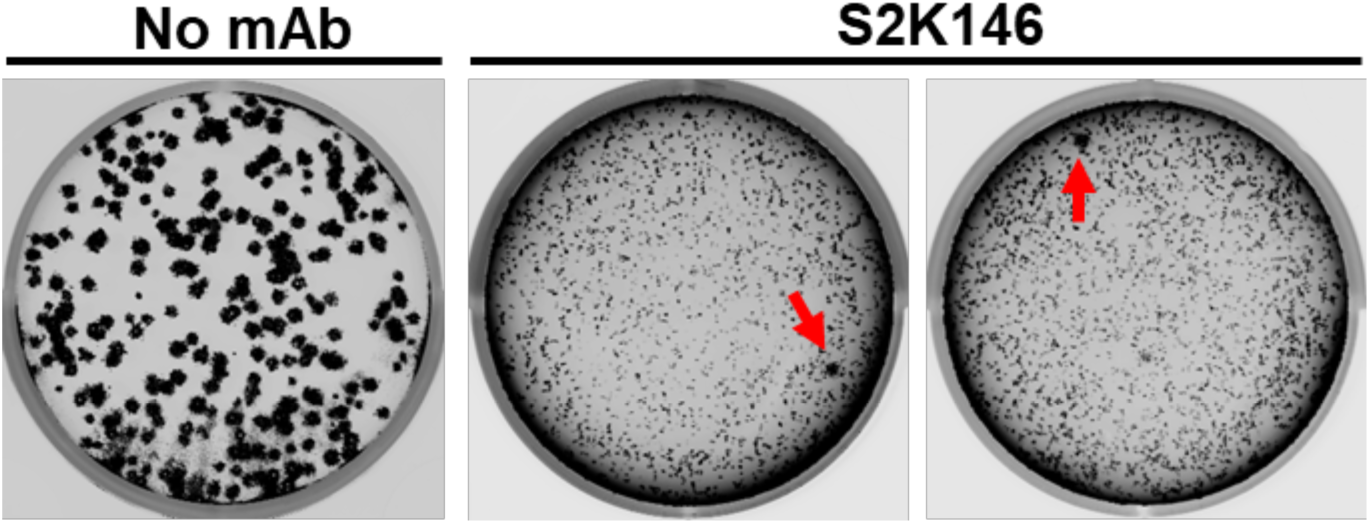
S2K146 escape clone selection by plaque assays using VSV-SARS-CoV-2 Wuhan-Hu-1 D614G S chimeric virus. Plaque assays performed using VSV-SARS-CoV-2 Wuhan-Hu-1 D614G S chimeric virus on Vero cells with or without S2K146 in the overlay to isolate escape mutants (red arrow). Data are representative of two sets of experiments.

**Figure S7.**
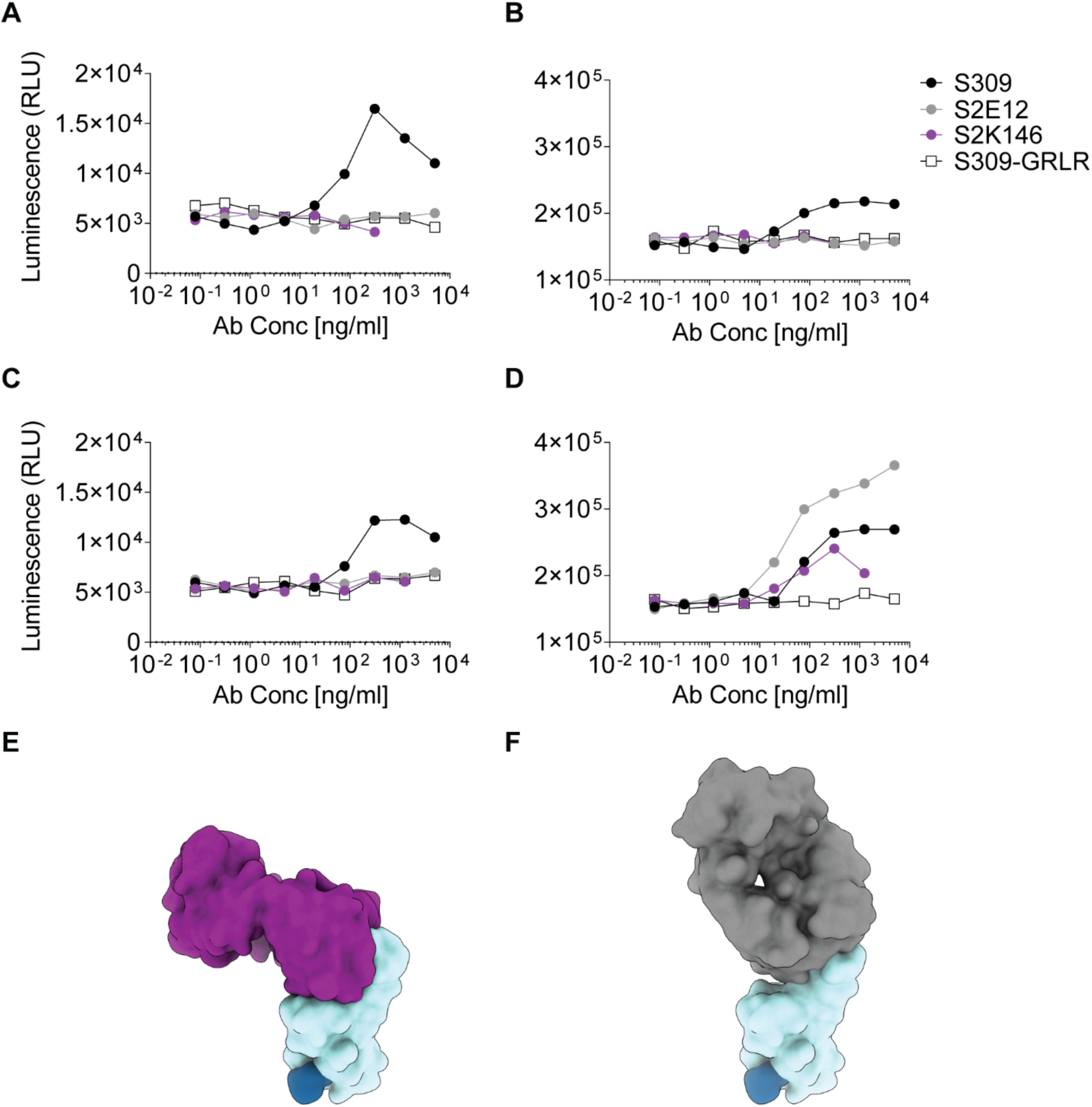
Activation of Fc**γ**RIIa AND Fc**γ**RllIa in vitro. **A-B)** NFAT-driven luciferase signal indued in Jurkat cells stably expressing Fc**γ**RIIa H131 variant (A) or Fc**γ**RIIIa V148 variant (B) by S2K146 mAb binding to full-length wild type SARS-CoV-2 S on CHO target cells. **C-D)** NFAT-driven luciferase signal induced in Jurkat cells stably expressing Fc**γ**Rlla H131 variant (C) or Fc**γ**RIIIa V148 variant (D) by S2K146 mAb binding to uncleavable full-length wild-type SARS-CoV-2 S on CHO target cells. **E)** Surface rendering of site-I-targeting S2K146 (purple) mAb bound to the SARS-CoV-2 RBD (light blue). **F)** Surface rendering of site-I-targeting S2E12 (grey) mAb bound to the SARS-CoV-2 RBD (light blue). The N343 glycan is rendered as a blue surface in E and F.

**Figure S8.**
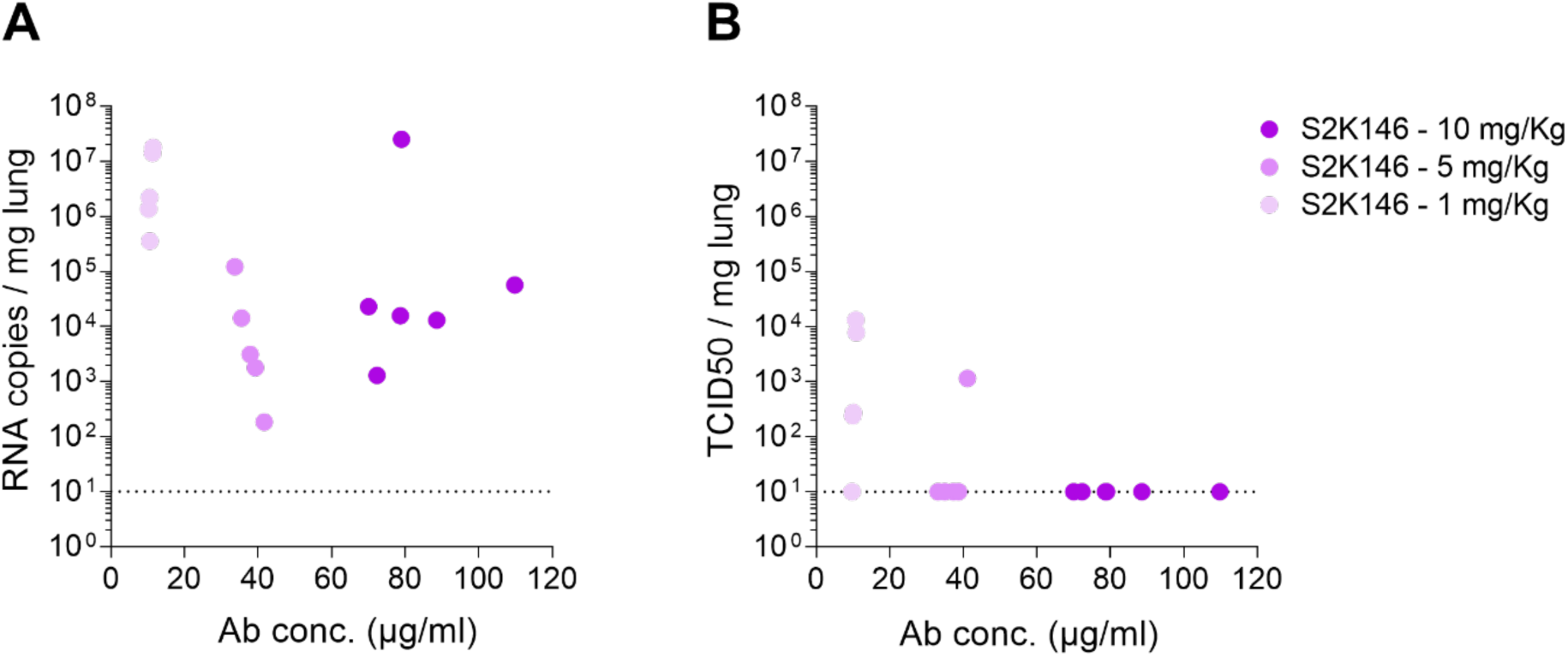
Correlation between mAb concentration and viral loads in the lungs of challenged hamsters. **(A-B)** Viral RNA copy number (A) and replicating virus titers (B) of SARS-CoV-2 Beta VOC in the lung of Syrian hamsters at 4 days post-infection plotted as a function of S2K146 mAb serum concentrations at day 1 post infection. (S2K146 10 mg/kg n = 6; S2K146 5 mg/kg n = 5; S2K146 5 mg/kg n = 5) was administered 1 day post infection.

**Table S1.** CryoEM data collection and refinement statistics.

|  | SARS-CoV-2<br>S/S2K146<br>(3RBDs open) | SARS-CoV-2<br>S/S2K146<br>(2RBDs<br>open) | SARS-CoV-2 S<br>RBD/S2K146<br>(Local refinement) |
| --- | --- | --- | --- |
| **Data collection**<br>**and processing** |  |  |  |
| Magnification | 105,000 | 105,000 | 105,000 |
| Voltage (kV) | 300 | 300 | 300 |
| Electron exposure<br>(e<sup>-</sup>/Å<sup>2</sup>) | 60 | 60 | 60 |
| Defocus range (μm) | 0.5-2.5 | 0.5-2.5 | 0.5-2.5 |
| Pixel size (Å) | 0.843 | 0.843 | 0.843 |
| Symmetry imposed | C1 | C1 | C1 |
| Final particle<br>images (no.) | 212,054 | 348,047 | 97,818 |
| Map resolution (Å) | 3.2 | 3.2 | 3.2 |
| FSC threshold | 0.143 | 0.143 | 0.143 |
| Map sharpening *B*<br>factor (Å<sup>2</sup>) | -96 | -99 | -52 |
| **Validation** |  |  |  |
| MolProbity score |  | 1.22 | 1.25 |
| Clashscore |  | 1.27 | 1.45 |
| Poor rotamers (%) |  | 0 | 0 |
| Ramachandran plot |  |  |  |
| Favored (%) |  | 95 | 96.2 |
| Allowed (%) |  | 4.55 | 3.55 |
| Disallowed (%) |  | 0.75 | 0.25 |

**Table S2.** Summary of nucleotide and amino acid mutations found in 36 neutralization-resistant VSV-SARS-CoV-2-S D614G chimera plaques.

| Clones | Nucleotide mutant | Amino acid Mutant |
| --- | --- | --- |
| #1 | T1465C/<br>T1602G | Y489H/<br>V534V |
| #2 | T1465C | Y489H |
| #3 | T1465C | Y489H |
| #4 | T1465C | Y489H |
| #5 | T1465C | Y489H |
| #6 | T1465C | Y489H |
| #7 | T1465C | Y489H |
| #8 | T1465C | Y489H |
| #9 | T1465C | Y489H |
| #10 | T1465C | Y489H |
| #11 | T1465C | Y489H |
| #12 | T1465C | Y489H |
| #13 | T1465C | Y489H |
| #14 | T1465C | Y489H |
| #15 | T1465C | Y489H |
| #16 | T1465C | Y489H |
| #17 | T1465C | Y489H |
| #18 | T1465C | Y489H |
| #19 | T1465C | Y489H |
| #20 | T1465C | Y489H |
| #21 | T1465C | Y489H |
| #22 | T1465C | Y489H |
| #23 | T1465C | Y489H |
| #24 | T1465C | Y489H |
| #25 | T1465C | Y489H |
| #26 | T1465C | Y489H |
| #27 | T1465C | Y489H |
| #28 | T1464C/<br>T1465C | C488C/<br>Y489H |
| #29 | T1465C | Y489H |
| #30 | T1263C/<br>T1465C | Y421Y/<br>Y489H |
| #31 | T1465C/<br>A1671G | Y489H/<br>K557K |
| #32 | T1465C | Y489H |
| #33 | T1465C | Y489H |
| #34 | T1465C | Y489H |
| #35 | T1465C | Y489H |
| #36 | T1465C | Y489H |

## References

1. C. A. Sánchez, H. Li, K. L. Phelps, C. Zambrana-Torrelio, L.-F. Wang, K. J. Olival, P. Daszak, A strategy to assess spillover risk of bat SARS-related coronaviruses in Southeast Asia. medRxiv (2021), doi:10.1101/2021.09.09.21263359.

2. A. C. Walls, M. A. Tortorici, B. J. Bosch, B. Frenz, P. J. M. Rottier, F. DiMaio, F. A. Rey, D. Veesler, Cryo-electron microscopy structure of a coronavirus spike glycoprotein trimer. Nature. 531, 114–117 (2016).

3. A. C. Walls, M. A. Tortorici, J. Snijder, X. Xiong, B. J. Bosch, F. A. Rey, D. Veesler, Tectonic conformational changes of a coronavirus spike glycoprotein promote membrane fusion. Proc. Natl. Acad. Sci. U. S. A. 114, 11157–11162 (2017).

4. A. C. Walls, Y. J. Park, M. A. Tortorici, A. Wall, A. T. McGuire, D. Veesler, Structure, Function, and Antigenicity of the SARS-CoV-2 Spike Glycoprotein. Cell. 181, 281–292.e6 (2020).

5. D. Wrapp, N. Wang, K. S. Corbett, J. A. Goldsmith, C. L. Hsieh, O. Abiona, B. S. Graham, J. S. McLellan, Cryo-EM structure of the 2019-nCoV spike in the prefusion conformation. Science. 367, 1260–1263 (2020).

6. M. A. Tortorici, D. Veesler, Structural insights into coronavirus entry. Adv. Virus Res. 105, 93–116 (2019).

7. P. Zhou, X. L. Yang, X. G. Wang, B. Hu, L. Zhang, W. Zhang, H. R. Si, Y. Zhu, B. Li, C. L. Huang, H. D. Chen, J. Chen, Y. Luo, H. Guo, R. D. Jiang, M. Q. Liu, Y. Chen, X. R. Shen, X. Wang, X. S. Zheng, K. Zhao, Q. J. Chen, F. Deng, L. L. Liu, B. Yan, F. X. Zhan, Y. Y. Wang, G. F. Xiao, Z. L. Shi, A pneumonia outbreak associated with a new coronavirus of probable bat origin. Nature (2020), doi:10.1038/s41586-020-2012-7.

8. M. Letko, A. Marzi, V. Munster, Functional assessment of cell entry and receptor usage for SARS-CoV-2 and other lineage B betacoronaviruses. Nature Microbiology (2020), doi:10.1038/s41564-020-0688-y.

9. M. Hoffmann, H. Kleine-Weber, S. Schroeder, N. Krüger, T. Herrler, S. Erichsen, T. S. Schiergens, G. Herrler, N. H. Wu, A. Nitsche, M. A. Müller, C. Drosten, S. Pöhlmann, SARS-CoV-2 Cell Entry Depends on ACE2 and TMPRSS2 and Is Blocked by a Clinically Proven Protease Inhibitor. Cell. 181, 271–280.e8 (2020).

10. W. Li, M. J. Moore, N. Vasilieva, J. Sui, S. K. Wong, M. A. Berne, M. Somasundaran, J. L. Sullivan, K. Luzuriaga, T. C. Greenough, H. Choe, M. Farzan, Angiotensin-converting enzyme 2 is a functional receptor for the SARS coronavirus. Nature. 426, 450–454 (2003).

11. L. Piccoli, Y. J. Park, M. A. Tortorici, N. Czudnochowski, A. C. Walls, M. Beltramello, C. Silacci-Fregni, D. Pinto, L. E. Rosen, J. E. Bowen, O. J. Acton, S. Jaconi, B. Guarino, A. Minola, F. Zatta, N. Sprugasci, J. Bassi, A. Peter, A. De Marco, J. C. Nix, F. Mele, S. Jovic, B. F. Rodriguez, S. V. Gupta, F. Jin, G. Piumatti, G. Lo Presti, A. F. Pellanda, M. Biggiogero, M. Tarkowski, M. S. Pizzuto, E. Cameroni, C. Havenar-Daughton, M. Smithey, D. Hong, V. Lepori, E. Albanese, A. Ceschi, E. Bernasconi, L. Elzi, P. Ferrari, C. Garzoni, A. Riva, G. Snell, F. Sallusto, K. Fink, H. W. Virgin, A. Lanzavecchia, D. Corti, D. Veesler, Mapping Neutralizing and Immunodominant Sites on the SARS-CoV-2 Spike Receptor-Binding Domain by Structure-Guided High-Resolution Serology. Cell. 183, 1024–1042.e21 (2020).

12. A. J. Greaney, A. N. Loes, L. E. Gentles, K. H. D. Crawford, T. N. Starr, K. D. Malone, H. Y. Chu, J. D. Bloom, Antibodies elicited by mRNA-1273 vaccination bind more broadly to the receptor binding domain than do those from SARS-CoV-2 infection. Sci. Transl. Med. 13 (2021), doi:10.1126/scitranslmed.abi9915.

13. M. A. Tortorici, N. Czudnochowski, T. N. Starr, R. Marzi, A. C. Walls, F. Zatta, J. E. Bowen, S. Jaconi, J. Di Iulio, Z. Wang, A. De Marco, S. K. Zepeda, D. Pinto, Z. Liu, M. Beltramello, I. Bartha, M. P. Housley, F. A. Lempp, L. E. Rosen, E. Dellota Jr, H. Kaiser, M. Montiel-Ruiz, J. Zhou, A. Addetia, B. Guarino, K. Culap, N. Sprugasci, C. Saliba, E. Vetti, I. Giacchetto-Sasselli, C. S. Fregni, R. Abdelnabi, S.-Y. C. Foo, C. Havenar-Daughton, M. A. Schmid, F. Benigni, E. Cameroni, J. Neyts, A. Telenti, H. W. Virgin, S. P. J. Whelan, G. Snell, J. D. Bloom, D. Corti, D. Veesler, M. S. Pizzuto, Broad sarbecovirus neutralization by a human monoclonal antibody. Nature (2021), doi:10.1038/s41586-021-03817-4.

14. T. N. Starr, N. Czudnochowski, Z. Liu, F. Zatta, Y.-J. Park, A. Addetia, D. Pinto, M. Beltramello, P. Hernandez, A. J. Greaney, R. Marzi, W. G. Glass, I. Zhang, A. S. Dingens, J. E. Bowen, M. A. Tortorici, A. C. Walls, J. A. Wojcechowskyj, A. De Marco, L. E. Rosen, J. Zhou, M. Montiel-Ruiz, H. Kaiser, J. Dillen, H. Tucker, J. Bassi, C. Silacci-Fregni, M. P. Housley, J. di Iulio, G. Lombardo, M. Agostini, N. Sprugasci, K. Culap, S. Jaconi, M. Meury, E. Dellota, R. Abdelnabi, S.-Y. C. Foo, E. Cameroni, S. Stumpf, T. I. Croll, J. C. Nix, C. Havenar-Daughton, L. Piccoli, F. Benigni, J. Neyts, A. Telenti, F. A. Lempp, M. S. Pizzuto, J. D. Chodera, C. M. Hebner, H. W. Virgin, S. P. J. Whelan, D. Veesler, D. Corti, J. D. Bloom, G. Snell, SARS-CoV-2 RBD antibodies that maximize breadth and resistance to escape. Nature (2021), doi:10.1038/s41586-021-03807-6.

15. D. Pinto, Y. J. Park, M. Beltramello, A. C. Walls, M. A. Tortorici, S. Bianchi, S. Jaconi, K. Culap, F. Zatta, A. De Marco, A. Peter, B. Guarino, R. Spreafico, E. Cameroni, J. B. Case, R. E. Chen, C. Havenar-Daughton, G. Snell, A. Telenti, H. W. Virgin, A. Lanzavecchia, M. S. Diamond, K. Fink, D. Veesler, D. Corti, Cross-neutralization of SARS-CoV-2 by a human monoclonal SARS-CoV antibody. Nature. 583, 290–295 (2020).

16. C. A. Jette, A. A. Cohen, P. N. P. Gnanapragasam, F. Muecksch, Y. E. Lee, K. E. Huey-Tubman, F. Schmidt, T. Hatziioannou, P. D. Bieniasz, M. C. Nussenzweig, A. P. West, J. R. Keeffe, P. J. Bjorkman, C. O. Barnes, Broad cross-reactivity across sarbecoviruses exhibited by a subset of COVID-19 donor-derived neutralizing antibodies. bioRxivorg (2021), doi:10.1101/2021.04.23.441195.

17. D. R. Martinez, A. Schaefer, S. Gobeil, D. Li, G. De la Cruz, R. Parks, X. Lu, M. Barr, K. Manne, K. Mansouri, R. J. Edwards, B. Yount, K. Anasti, S. A. Montgomery, S. Shen, T. Zhou, P. D. Kwong, B. S. Graham, J. R. Mascola, D. C. Montefiori, M. Alam, G. D. Sempowski, K. Wiehe, K. O. Saunders, P. Acharya, B. F. Haynes, R. S. Baric, A broadly neutralizing antibody protects against SARS-CoV, pre-emergent bat CoVs, and SARS-CoV-2 variants in mice. bioRxivorg (2021), doi:10.1101/2021.04.27.441655.

18. A. Z. Wec, D. Wrapp, A. S. Herbert, D. P. Maurer, D. Haslwanter, M. Sakharkar, R. K. Jangra, M. E. Dieterle, A. Lilov, D. Huang, L. V. Tse, N. V. Johnson, C. L. Hsieh, N. Wang, J. H. Nett, E. Champney, I. Burnina, M. Brown, S. Lin, M. Sinclair, C. Johnson, S. Pudi, R. Bortz, A. S. Wirchnianski, E. Laudermilch, C. Florez, J. M. Fels, C. M. O’Brien, B. S. Graham, D. Nemazee, D. R. Burton, R. S. Baric, J. E. Voss, K. Chandran, J. M. Dye, J. S. McLellan, L. M. Walker, Broad neutralization of SARS-related viruses by human monoclonal antibodies. Science (2020), doi:10.1126/science.abc7424.

19. C. G. Rappazzo, L. V. Tse, C. I. Kaku, D. Wrapp, M. Sakharkar, D. Huang, L. M. Deveau, T. J. Yockachonis, A. S. Herbert, M. B. Battles, C. M. O’Brien, M. E. Brown, J. C. Geoghegan, J. Belk, L. Peng, L. Yang, Y. Hou, T. D. Scobey, D. R. Burton, D. Nemazee, J. M. Dye, J. E. Voss, B. M. Gunn, J. S. McLellan, R. S. Baric, L. E. Gralinski, L. M. Walker, Broad and potent activity against SARS-like viruses by an engineered human monoclonal antibody. Science. 371, 823–829 (2021).

20. D. Corti, L. A. Purcell, G. Snell, D. Veesler, Tackling COVID-19 with neutralizing monoclonal antibodies. Cell (2021), doi:10.1016/j.cell.2021.05.005.

21. M. McCallum, J. Bassi, A. De Marco, A. Chen, A. C. Walls, J. Di Iulio, M. A. Tortorici, M.-J. Navarro, C. Silacci-Fregni, C. Saliba, K. R. Sprouse, M. Agostini, D. Pinto, K. Culap, S. Bianchi, S. Jaconi, E. Cameroni, J. E. Bowen, S. W. Tilles, M. S. Pizzuto, S. B. Guastalla, G. Bona, A. F. Pellanda, C. Garzoni, W. C. Van Voorhis, L. E. Rosen, G. Snell, A. Telenti, H. W. Virgin, L. Piccoli, D. Corti, D. Veesler, SARS-CoV-2 immune evasion by the B.1.427/B.1.429 variant of concern. Science (2021), doi:10.1126/science.abi7994.

22. M. McCallum, A. C. Walls, K. R. Sprouse, J. E. Bowen, L. Rosen, H. V. Dang, A. deMarco, N. Franko, S. W. Tilles, J. Logue, M. C. Miranda, M. Ahlrichs, L. Carter, G. Snell, M. S. Pizzuto, H. Y. Chu, W. C. Van Voorhis, D. Corti, D. Veesler, Molecular basis of immune evasion by the delta and kappa SARS-CoV-2 variants. bioRxivorg (2021), doi:10.1101/2021.08.11.455956.

23. P. Mlcochova, S. Kemp, M. S. Dhar, G. Papa, B. Meng, I. A. T. M. Ferreira, R. Datir, D. A. Collier, A. Albecka, S. Singh, R. Pandey, J. Brown, J. Zhou, N. Goonawardane, S. Mishra, C. Whittaker, T. Mellan, R. Marwal, M. Datta, S. Sengupta, K. Ponnusamy, V. S. Radhakrishnan, A. Abdullahi, O. Charles, P. Chattopadhyay, P. Devi, D. Caputo, T. Peacock, D. C. Wattal, N. Goel, A. Satwik, R. Vaishya, M. Agarwal, Indian SARS-CoV-2 Genomics Consortium (INSACOG), Genotype to Phenotype Japan (G2P-Japan) Consortium, CITIID-NIHR BioResource COVID-19 Collaboration, A. Mavousian, J. H. Lee, J. Bassi, C. Silacci-Fegni, C. Saliba, D. Pinto, T. Irie, I. Yoshida, W. L. Hamilton, K. Sato, S. Bhatt, S. Flaxman, L. C. James, D. Corti, L. Piccoli, W. S. Barclay, P. Rakshit, A. Agrawal, R. K. Gupta, SARS-CoV-2 B.1.617.2 Delta variant replication and immune evasion. Nature (2021), doi:10.1038/s41586-021-03944-y.

24. T. N. Starr, A. J. Greaney, S. K. Hilton, D. Ellis, K. H. D. Crawford, A. S. Dingens, M. J. Navarro, J. E. Bowen, M. A. Tortorici, A. C. Walls, N. P. King, D. Veesler, J. D. Bloom, Deep Mutational Scanning of SARS-CoV-2 Receptor Binding Domain Reveals Constraints on Folding and ACE2 Binding. Cell. 182, 1295–1310.e20 (2020).

25. T. N. Starr, A. J. Greaney, A. S. Dingens, J. D. Bloom, Complete map of SARS-CoV-2 RBD mutations that escape the monoclonal antibody LY-CoV555 and its cocktail with LY-CoV016. Cell Rep Med. 2, 100255 (2021).

26. T. N. Starr, A. J. Greaney, A. Addetia, W. W. Hannon, M. C. Choudhary, A. S. Dingens, J. Z. Li, J. D. Bloom, Prospective mapping of viral mutations that escape antibodies used to treat COVID-19. Science. 371, 850–854 (2021).

27. M. A. Tortorici, M. Beltramello, F. A. Lempp, D. Pinto, H. V. Dang, L. E. Rosen, M. McCallum, J. Bowen, A. Minola, S. Jaconi, F. Zatta, A. De Marco, B. Guarino, S. Bianchi, E. J. Lauron, H. Tucker, J. Zhou, A. Peter, C. Havenar-Daughton, J. A. Wojcechowskyj, J. B. Case, R. E. Chen, H. Kaiser, M. Montiel-Ruiz, M. Meury, N. Czudnochowski, R. Spreafico, J. Dillen, C. Ng, N. Sprugasci, K. Culap, F. Benigni, R. Abdelnabi, S. C. Foo, M. A. Schmid, E. Cameroni, Riva, A. Gabrieli, M. Galli, M. S. Pizzuto, J. Neyts, M. S. Diamond, H. W. Virgin, G. Snell, D. Corti, K. Fink, D. Veesler, Ultrapotent human antibodies protect against SARS-CoV-2 challenge via multiple mechanisms. Science. 370, 950–957 (2020).

28. T. N. Starr, S. K. Zepeda, A. C. Walls, A. J. Greaney, D. Veesler, J. D. Bloom, ACE2 binding is an ancestral and evolvable trait of sarbecoviruses. bioRxiv (2021), doi:10.1101/2021.07.17.452804.

29. F. Li, W. Li, M. Farzan, S. C. Harrison, Structure of SARS coronavirus spike receptor-binding domain complexed with receptor. Science. 309, 1864–1868 (2005).

30. J. B. Case, P. W. Rothlauf, R. E. Chen, Z. Liu, H. Zhao, A. S. Kim, L. M. Bloyet, Q. Zeng, S. Tahan, L. Droit, M. X. G. Ilagan, M. A. Tartell, G. Amarasinghe, J. P. Henderson, S. Miersch, M. Ustav, S. Sidhu, H. W. Virgin, D. Wang, S. Ding, D. Corti, E. S. Theel, D. H. Fremont, M. S. Diamond, S. P. J. Whelan, Neutralizing Antibody and Soluble ACE2 Inhibition of a Replication-Competent VSV-SARS-CoV-2 and a Clinical Isolate of SARS-CoV-2. Cell Host Microbe. 28, 475–485.e5 (2020).

31. A. C. Walls, X. Xiong, Y. J. Park, M. A. Tortorici, J. Snijder, J. Quispe, E. Cameroni, R. Gopal, M. Dai, A. Lanzavecchia, M. Zambon, F. A. Rey, D. Corti, D. Veesler, Unexpected Receptor Functional Mimicry Elucidates Activation of Coronavirus Fusion. Cell. 176, 1026–1039.e15 (2019).

32. F. A. Lempp, L. Soriaga, M. Montiel-Ruiz, F. Benigni, J. Noack, Y.-J. Park, S. Bianchi, A. C. Walls, J. E. Bowen, J. Zhou, H. Kaiser, A. Joshi, M. Agostini, M. Meury, E. Dellota Jr, S. Jaconi, E. Cameroni, J. Martinez-Picado, J. Vergara-Alert, N. Izquierdo-Useros, H. W. Virgin, A. Lanzavecchia, D. Veesler, L. Purcell, A. Telenti, D. Corti, Lectins enhance SARS-CoV-2 infection and influence neutralizing antibodies. Nature (2021), doi:10.1038/s41586-021-03925-1.

33. J. Huo, Y. Zhao, J. Ren, D. Zhou, H. M. E. Duyvesteyn, H. M. Ginn, L. Carrique, T. Malinauskas, R. R. Ruza, P. N. M. Shah, T. K. Tan, P. Rijal, N. Coombes, K. R. Bewley, J. A. Tree, J. Radecke, N. G. Paterson, P. Supasa, J. Mongkolsapaya, G. R. Screaton, M. Carroll, A. Townsend, E. E. Fry, R. J. Owens, D. I. Stuart, Neutralisation of SARS-CoV-2 by destruction of the prefusion Spike. Cell Host Microbe (2020), doi:10.1016/j.chom.2020.06.010.

34. R. Abdelnabi, R. Boudewijns, C. S. Foo, L. Seldeslachts, L. Sanchez-Felipe, X. Zhang, L. Delang, P. Maes, S. J. F. Kaptein, B. Weynand, G. V. Velde, J. Neyts, K. Dallmeier, Comparative infectivity and pathogenesis of emerging SARS-CoV-2 variants in Syrian hamsters (2021), doi:10.1101/2021.02.26.433062.

35. R. Boudewijns, H. J. Thibaut, S. J. F. Kaptein, R. Li, V. Vergote, L. Seldeslachts, J. Van Weyenbergh, C. De Keyzer, L. Bervoets, S. Sharma, L. Liesenborghs, J. Ma, S. Jansen, D. Van Looveren, T. Vercruysse, X. Wang, D. Jochmans, E. Martens, K. Roose, D. De Vlieger, B. Schepens, T. Van Buyten, S. Jacobs, Y. Liu, J. Martí-Carreras, B. Vanmechelen, T. Wawina-Bokalanga, L. Delang, J. Rocha-Pereira, L. Coelmont, W. Chiu, P. Leyssen, E. Heylen, D. Schols, L. Wang, L. Close, J. Matthijnssens, M. Van Ranst, V. Compernolle, G. Schramm, K. Van Laere, X. Saelens, N. Callewaert, G. Opdenakker, P. Maes, B. Weynand, C. Cawthorne, G. Vande Velde, Z. Wang, J. Neyts, K. Dallmeier, STAT2 signaling restricts viral dissemination but drives severe pneumonia in SARS-CoV-2 infected hamsters. Nat. Commun. 11, 5838 (2020).

36. J. Lan, J. Ge, J. Yu, S. Shan, H. Zhou, S. Fan, Q. Zhang, X. Shi, Q. Wang, L. Zhang, X. Wang, Structure of the SARS-CoV-2 spike receptor-binding domain bound to the ACE2 receptor. Nature (2020), doi:10.1038/s41586-020-2180-5.

37. W. Dejnirattisai, D. Zhou, H. M. Ginn, H. M. E. Duyvesteyn, P. Supasa, J. B. Case, Y. Zhao, T. S. Walter, A. J. Mentzer, C. Liu, B. Wang, G. C. Paesen, J. Slon-Campos, C. López-Camacho, N. M. Kafai, A. L. Bailey, R. E. Chen, B. Ying, C. Thompson, J. Bolton, A. Fyfe, S. Gupta, T. K. Tan, J. Gilbert-Jaramillo, W. James, M. Knight, M. W. Carroll, D. Skelly, C. Dold, Y. Peng, R. Levin, T. Dong, A. J. Pollard, J. C. Knight, P. Klenerman, N. Temperton, D. R. Hall, M. A. Williams, N. G. Paterson, F. K. R. Bertram, C. A. Seibert, D. K. Clare, A. Howe, J. Raedecke, Y. Song, A. R. Townsend, K.-Y. A. Huang, E. E. Fry, J. Mongkolsapaya, M. S. Diamond, J. Ren, D. I. Stuart, G. R. Screaton, The antigenic anatomy of SARS-CoV-2 receptor binding domain. Cell (2021), doi:10.1016/j.cell.2021.02.032.

38. P. S. Arunachalam, A. C. Walls, N. Golden, C. Atyeo, S. Fischinger, C. Li, P. Aye, M. J. Navarro, L. Lai, V. V. Edara, K. Röltgen, K. Rogers, L. Shirreff, D. E. Ferrell, S. Wrenn, D. Pettie, J. C. Kraft, M. C. Miranda, E. Kepl, C. Sydeman, N. Brunette, M. Murphy, B. Fiala, L. Carter, A. G. White, M. Trisal, C.-L. Hsieh, K. Russell-Lodrigue, C. Monjure, J. Dufour, S. Spencer, L. Doyle-Meyer, R. P. Bohm, N. J. Maness, C. Roy, J. A. Plante, K. S. Plante, A. Zhu, M. J. Gorman, S. Shin, X. Shen, J. Fontenot, S. Gupta, D. T. O’Hagan, R. Van Der Most, R. Rappuoli, R. L. Coffman, D. Novack, J. S. McLellan, S. Subramaniam, D. Montefiori, S. D. Boyd, J. L. Flynn, G. Alter, F. Villinger, H. Kleanthous, J. Rappaport, M. S. Suthar, N. P. King, D. Veesler, B. Pulendran, Adjuvanting a subunit COVID-19 vaccine to induce protective immunity. Nature (2021), doi:10.1038/s41586-021-03530-2.

39. A. C. Walls, B. Fiala, A. Schäfer, S. Wrenn, M. N. Pham, M. Murphy, L. V. Tse, L. Shehata, M. A. O’Connor, C. Chen, M. J. Navarro, M. C. Miranda, D. Pettie, R. Ravichandran, J. C. Kraft, C. Ogohara, A. Palser, S. Chalk, E. C. Lee, K. Guerriero, E. Kepl, C. M. Chow, C. Sydeman, E. A. Hodge, B. Brown, J. T. Fuller, K. H. Dinnon, L. E. Gralinski, S. R. Leist, K. L. Gully, T. B. Lewis, M. Guttman, H. Y. Chu, K. K. Lee, D. H. Fuller, R. S. Baric, P. Kellam, L. Carter, M. Pepper, T. P. Sheahan, D. Veesler, N. P. King, Elicitation of Potent Neutralizing Antibody Responses by Designed Protein Nanoparticle Vaccines for SARS-CoV-2. Cell. 183, 1367–1382.e17 (2020).

40. A. C. Walls, M. C. Miranda, A. Schäfer, M. N. Pham, A. Greaney, P. S. Arunachalam, M.-J. Navarro, M. A. Tortorici, K. Rogers, M. A. O’Connor, L. Shirreff, D. E. Ferrell, J. Bowen, N. Brunette, E. Kepl, S. K. Zepeda, T. Starr, C.-L. Hsieh, B. Fiala, S. Wrenn, D. Pettie, C. Sydeman, K. R. Sprouse, M. Johnson, A. Blackstone, R. Ravichandran, C. Ogohara, L. Carter, S. W. Tilles, R. Rappuoli, S. R. Leist, D. R. Martinez, M. Clark, R. Tisch, D. T. O’Hagan, R. Van Der Most, W. C. Van Voorhis, D. Corti, J. S. McLellan, H. Kleanthous, T. P. Sheahan, K. D. Smith, D. H. Fuller, F. Villinger, J. Bloom, B. Pulendran, R. Baric, N. P. King, D. Veesler, Elicitation of broadly protective sarbecovirus immunity by receptor-binding domain nanoparticle vaccines. Cell (2021), doi:10.1016/j.cell.2021.09.015.

41. K. O. Saunders, E. Lee, R. Parks, D. R. Martinez, D. Li, H. Chen, R. J. Edwards, S. Gobeil, M. Barr, K. Mansouri, S. M. Alam, L. L. Sutherland, F. Cai, A. M. Sanzone, M. Berry, K. Manne, K. W. Bock, M. Minai, B. M. Nagata, A. B. Kapingidza, M. Azoitei, L. V. Tse, T. D. Scobey, R. L. Spreng, R. W. Rountree, C. T. DeMarco, T. N. Denny, C. W. Woods, E. W. Petzold, J. Tang, T. H. Oguin 3rd, G. D. Sempowski, M. Gagne, D. C. Douek, M. A. Tomai, C. B. Fox, R. Seder, K. Wiehe, D. Weissman, N. Pardi, H. Golding, S. Khurana, P. Acharya, H. Andersen, M. G. Lewis, I. N. Moore, D. C. Montefiori, R. S. Baric, B. F. Haynes, Neutralizing antibody vaccine for pandemic and pre-emergent coronaviruses. Nature. 594, 553–559 (2021).

42. D. R. Martinez, A. Schäfer, S. R. Leist, G. De la Cruz, A. West, E. N. Atochina-Vasserman, L. C. Lindesmith, N. Pardi, R. Parks, M. Barr, D. Li, B. Yount, K. O. Saunders, D. Weissman, B. F. Haynes, S. A. Montgomery, R. S. Baric, Chimeric spike mRNA vaccines protect against Sarbecovirus challenge in mice. Science (2021), doi:10.1126/science.abi4506.

43. D. Pinto, M. M. Sauer, N. Czudnochowski, J. S. Low, M. Alejandra Tortorici, M. P. Housley, J. Noack, A. C. Walls, J. E. Bowen, B. Guarino, L. E. Rosen, J. di Iulio, J. Jerak, H. Kaiser, S. Islam, S. Jaconi, N. Sprugasci, K. Culap, R. Abdelnabi, C. Foo, L. Coelmont, I. Bartha, S. Bianchi, C. Silacci-Fregni, J. Bassi, R. Marzi, E. Vetti, A. Cassotta, A. Ceschi, P. Ferrari, P. E. Cippà, O. Giannini, S. Ceruti, C. Garzoni, A. Riva, F. Benigni, E. Cameroni, L. Piccoli, M. S. Pizzuto, M. Smithey, D. Hong, A. Telenti, F. A. Lempp, J. Neyts, C. Havenar-Daughton, A. Lanzavecchia, F. Sallusto, G. Snell, H. W. Virgin, M. Beltramello, D. Corti, D. Veesler, Broad betacoronavirus neutralization by a stem helix–specific human antibody. Science (2021), doi:10.1126/science.abj3321.

44. M. M. Sauer, M. A. Tortorici, Y.-J. Park, A. C. Walls, L. Homad, O. J. Acton, J. E. Bowen, C. Wang, X. Xiong, W. de van der Schueren, J. Quispe, B. G. Hoffstrom, B.-J. Bosch, A. T. McGuire, D. Veesler, Structural basis for broad coronavirus neutralization. Nat. Struct. Mol. Biol. 28, 478–486 (2021).

45. G. Song, W.-T. He, S. Callaghan, F. Anzanello, D. Huang, J. Ricketts, J. L. Torres, N. Beutler, L. Peng, S. Vargas, J. Cassell, M. Parren, L. Yang, C. Ignacio, D. M. Smith, J. E. Voss, D. Nemazee, A. B. Ward, T. Rogers, D. R. Burton, R. Andrabi, Cross-reactive serum and memory B-cell responses to spike protein in SARS-CoV-2 and endemic coronavirus infection. Nat. Commun. 12, 2938 (2021).

46. P. Zhou, M. Yuan, G. Song, N. Beutler, N. Shaabani, D. Huang, W.-T. He, X. Zhu, S. Callaghan, P. Yong, F. Anzanello, L. Peng, J. Ricketts, M. Parren, E. Garcia, S. A. Rawlings, D. M. Smith, D. Nemazee, J. R. Teijaro, T. F. Rogers, I. A. Wilson, D. R. Burton, R. Andrabi, A protective broadly cross-reactive human antibody defines a conserved site of vulnerability on beta-coronavirus spikes. bioRxivorg (2021), doi:10.1101/2021.03.30.437769.

47. C. Wang, R. van Haperen, J. Gutiérrez-Álvarez, W. Li, N. M. A. Okba, I. Albulescu, I. Widjaja, B. van Dieren, R. Fernandez-Delgado, I. Sola, D. L. Hurdiss, O. Daramola, F. Grosveld, F. J. M. van Kuppeveld, B. L. Haagmans, L. Enjuanes, D. Drabek, B.-J. Bosch, A conserved immunogenic and vulnerable site on the coronavirus spike protein delineated by cross-reactive monoclonal antibodies. Nat. Commun. 12, 1715 (2021).

48. A. L. Cathcart, C. Havenar-Daughton, F. A. Lempp, D. Ma, M. Schmid, M. L. Agostini, B. Guarino, J. Di iulio, L. Rosen, H. Tucker, J. Dillen, S. Subramanian, B. Sloan, S. Bianchi, J. Wojcechowskyj, J. Zhou, H. Kaiser, A. Chase, M. Montiel-Ruiz, N. Czudnochowski, E. Cameroni, S. Ledoux, C. Colas, L. Soriaga, A. Telenti, S. Hwang, G. Snell, H. W. Virgin, D. Corti, C. M. Hebner, The dual function monoclonal antibodies VIR-7831 and VIR-7832 demonstrate potent in vitro and in vivo activity against SARS-CoV-2. bioRxiv (2021),, doi:10.1101/2021.03.09.434607.

49. C. L. Hsieh, J. A. Goldsmith, J. M. Schaub, A. M. DiVenere, H. C. Kuo, K. Javanmardi, K. C. Le, D. Wrapp, A. G. Lee, Y. Liu, C. W. Chou, P. O. Byrne, C. K. Hjorth, N. V. Johnson, J. Ludes-Meyers, A. W. Nguyen, J. Park, N. Wang, D. Amengor, J. J. Lavinder, G. C. Ippolito, J. A. Maynard, I. J. Finkelstein, J. S. McLellan, Structure-based design of prefusion-stabilized SARS-CoV-2 spikes. Science (2020), doi:10.1126/science.abd0826.

50. D. Pinto, C. Fenwick, C. Caillat, C. Silacci, S. Guseva, F. Dehez, C. Chipot, S. Barbieri, A. Minola, D. Jarrossay, G. D. Tomaras, X. Shen, A. Riva, M. Tarkowski, O. Schwartz, T. Bruel, J. Dufloo, M. S. Seaman, D. C. Montefiori, A. Lanzavecchia, D. Corti, G. Pantaleo, W. Weissenhorn, Structural basis for broad HIV-1 neutralization by the MPER-specific human broadly neutralizing antibody LN01. Cell Host Microbe. 26, 623–637.e8 (2019).

51. Y. Kaname, H. Tani, C. Kataoka, M. Shiokawa, S. Taguwa, T. Abe, K. Moriishi, T. Kinoshita, Y. Matsuura, Acquisition of complement resistance through incorporation of CD55/decay-accelerating factor into viral particles bearing baculovirus GP64. J. Virol. 84, 3210–3219 (2010).

52. A. J. Greaney, T. N. Starr, P. Gilchuk, S. J. Zost, E. Binshtein, A. N. Loes, S. K. Hilton, J. Huddleston, R. Eguia, K. H. D. Crawford, A. S. Dingens, R. S. Nargi, R. E. Sutton, N. Suryadevara, P. W. Rothlauf, Z. Liu, S. P. J. Whelan, R. H. Carnahan, J. E. Crowe, J. D. Bloom, Complete Mapping of Mutations to the SARS-CoV-2 Spike Receptor-Binding Domain that Escape Antibody Recognition. Cell Host Microbe (2020), doi:10.1016/j.chom.2020.11.007.

53. C. Suloway, J. Pulokas, D. Fellmann, A. Cheng, F. Guerra, J. Quispe, S. Stagg, C. S. Potter, B. Carragher, Automated molecular microscopy: the new Leginon system. J. Struct. Biol. 151, 41–60 (2005).

54. D. Tegunov, P. Cramer, Real-time cryo-electron microscopy data preprocessing with Warp. Nat. Methods. 16, 1146–1152 (2019).

55. A. Punjani, J. L. Rubinstein, D. J. Fleet, M. A. Brubaker, cryoSPARC: algorithms for rapid unsupervised cryo-EM structure determination. Nat. Methods. 14, 290–296 (2017).

56. J. Zivanov, T. Nakane, B. O. Forsberg, D. Kimanius, W. J. Hagen, E. Lindahl, S. H. Scheres, New tools for automated high-resolution cryo-EM structure determination in RELION-3. Elife. 7 (2018), doi:10.7554/eLife.42166.

57. A. Punjani, H. Zhang, D. J. Fleet, Non-uniform refinement: adaptive regularization improves single-particle cryo-EM reconstruction. Nat. Methods. 17, 1214–1221 (2020).

58. J. Zivanov, T. Nakane, S. H. W. Scheres, A Bayesian approach to beam-induced motion correction in cryo-EM single-particle analysis. IUCrJ. 6, 5–17 (2019).

59. S. Chen, G. McMullan, A. R. Faruqi, G. N. Murshudov, J. M. Short, S. H. Scheres, R. Henderson, High-resolution noise substitution to measure overfitting and validate resolution in 3D structure determination by single particle electron cryomicroscopy. Ultramicroscopy. 135, 24–35 (2013).

60. P. B. Rosenthal, R. Henderson, Optimal determination of particle orientation, absolute hand, and contrast loss in single-particle electron cryomicroscopy. J. Mol. Biol. 333, 721–745 (2003).

61. E. F. Pettersen, T. D. Goddard, C. C. Huang, G. S. Couch, D. M. Greenblatt, E. C. Meng, T. E. Ferrin, UCSF Chimera--a visualization system for exploratory research and analysis. J. Comput. Chem. 25, 1605–1612 (2004).

62. P. Emsley, B. Lohkamp, W. G. Scott, K. Cowtan, Features and development of Coot. Acta Crystallogr. D Biol. Crystallogr. 66, 486–501 (2010).

63. B. Frenz, S. Rämisch, A. J. Borst, A. C. Walls, J. Adolf-Bryfogle, W. R. Schief, D. Veesler, F. DiMaio, Automatically Fixing Errors in Glycoprotein Structures with Rosetta. Structure. 27, 134–139.e3 (2019).

64. R. Y. Wang, Y. Song, B. A. Barad, Y. Cheng, J. S. Fraser, F. DiMaio, Automated structure refinement of macromolecular assemblies from cryo-EM maps using Rosetta. Elife. 5 (2016), doi:10.7554/eLife.17219.

65. R. Abdelnabi, R. Boudewijns, C. S. Foo, L. Seldeslachts, L. Sanchez-Felipe, X. Zhang, L. Delang, P. Maes, S. J. F. Kaptein, B. Weynand, G. Vande Velde, J. Neyts, K. Dallmeier, Comparing infectivity and virulence of emerging SARS-CoV-2 variants in Syrian hamsters. EBioMedicine. 68, 103403 (2021).

66. L. J. Reed, H. Muench, A SIMPLE METHOD OF ESTIMATING FIFTY PER CENT ENDPOINTS12. Am. J. Epidemiol. 27, 493–497 (1938).

